# A MET-Targeted Variable New Antigen Receptor (VNAR) Theranostic for Non-Small Cell Lung Cancer

**DOI:** 10.64898/2026.01.30.702875

**Authors:** Rachel L. Minne, Jayden L. West, Natalie Y. Luo, Kwangok P. Nickel, Gihan S. Gunaratne, Kendahl L. Ott, Joseph P. Gallant, Kendall E. Barrett, Caroline M. Mork, Saahil Javeri, Madalynn R. Wopat, Loren D. Rivera Lopez, Will A. Toscano, Nina C. Zitzer, Ohyun Kwon, Jordan Teague, Brittany Bunker, Jessica M. Phillips, Malick Bio Idrissou, Hansel Comas Rojas, Jason C. Mixdorf, Eduardo Aluicio-Sarduy, Jonathan W. Engle, Bryan Bednarz, Reinier Hernandez, Randall J. Kimple, Andrew M. Baschnagel, Aaron M. LeBeau

## Abstract

The MET receptor tyrosine kinase is mutated or amplified in ∼6% of non-small cell lung cancer (NSCLC) and overexpressed in ∼80% of all NSCLC cases. A theranostic agent that can both see and treat MET-altered NSCLC has never been described before in the literature. Here, we report a shark-derived single-domain variable new antigen receptor (VNAR) for MET with theranostic applications. Following the immunization of a juvenile nurse shark (*Ginglymostoma cirratum*) with the extracellular domain of human MET, we identified a VNAR clone that specifically engaged MET with high affinity. Engineering the lead VNAR into a bivalent human Fc, vMET1-Fc, yielded a construct that selectively targeted and was internalized by MET-positive cells without affecting cell viability or downstream MET signaling. When radiolabeled with the positron emitting isotope Zr-89, [^89^Zr]Zr-vMET1-Fc enabled longitudinal PET/CT imaging. High tumor uptake with low background was observed in MET-positive NSCLC xenografts administered [^89^Zr]Zr-vMET1-Fc. As a targeted beta-particle radiotherapy, [¹⁷⁷Lu]Lu-vMET1-Fc resulted in marked tumor-growth delay and exhibited a favorable toxicity profile, collectively improving progression-free survival in NSCLC mouse models. Non-human primate PET/CT imaging studies with ([⁸⁹Zr]Zr-vMET1-Fc in healthy rhesus macaques confirmed favorable biodistribution and dosimetry, predictable clearance, and minimal off-target uptake. Additional blood chemistry analysis found no significant immune response or cytotoxicity. Together, these findings establish vMET1-Fc as a theranostic agent for imaging and treating MET-altered NSCLC.

**Statement of Significance:** A shark-derived antibody selectively targeting MET shows preclinical efficacy as a theranostic agent for MET-altered cancer.

## Introduction

MET, also known as c-MET, is a receptor tyrosine kinase that regulates cell proliferation, embryogenesis, and wound healing (1, 2). Dysregulation of MET, through amplification or mutation, drives oncogenesis by promoting unchecked cell growth and metastasis (3–6). In non-small cell lung cancer (NSCLC), MET is amplified in 3-6% (7), overexpressed in 30% (8), and mutated in 3-4% (9), with MET exon 14 skipping (METex14) being the most common mutation (9–11). METex14 mutations occur in the juxtamembrane domain, causing loss of the CBL binding site, impairing receptor degradation, and increasing MET expression on the cell surface (12). These mutations typically occur in the absence of other driver mutations and are associated with a poor prognosis (12, 14, 15). Selective tyrosine kinase inhibitors (TKIs) capmatinib and tepotinib have demonstrated clinical efficacy with response rates of 50-68% and are now used as first-line therapy in patients with METex14 NSCLC (16, 17). Despite these advances, resistance inevitably develops underscoring the need for new therapeutic strategies. Given the high surface expression of MET in both METex14-mutated and MET-amplified NSCLC, MET-directed antibodies are now being evaluated as promising therapeutic modalities.

Monoclonal antibodies represent a broad class of targeted therapeutics used across many disease settings. Conventional human immunoglobulin (IgG) molecules, which are composed of paired heavy and light chains, have historically served as the dominant antibody format. However, these traditional IgG antibodies are limited by suboptimal pharmacokinetic behavior and high manufacturing costs (17). Additionally, the structural organization and size of the complementarity determining regions (CDRs) constrains the flexibility with which IgGs recognize antigens, leading to a bias toward relatively flat epitopes and narrowing the range of molecular interactions they can form (18–20). Such constraints may have contributed to the inconsistent clinical performance of some antibody-based therapeutics, including those directed at MET (22, 23). Currently, MET-directed antibody therapies in NSCLC are in active clinical development; for example, the antibody-drug conjugate telisotuzumab vedotin (Teliso-V) is utilized for patients with high MET expression, while bispecific antibodies like amivantamab continue to be evaluated in combination regimens (24, 25). However, these therapies remain limited by the structural constraints of the traditional IgG format. The limitations of conventional antibodies have spurred interest in alternative molecular formats that offer greater design flexibility, including single-domain binding scaffolds (25–27). Among these, camelid-derived single domain antibodies (VHH) have received the most extensive characterization (25–27). In the last six years, VHH-derived therapies have been approved by the FDA and equivalent international agencies for conditions ranging from blood disorders (28) to rheumatoid arthritis (29). Our prior work has shown that MET-directed camelid antibodies can be used as molecular imaging agents for MET-expressing cancers (31, 32).

Less studied as targeting vectors are the single-domain binding regions isolated from IgNAR antibodies of elasmobranchs (sharks, rays, skates). These domains, called variable new antigen receptors (VNARs), have a molecular weight of 11-12 kDa, and are smaller than the equivalent camelid domains (15 kDa) and human single-chain variable fragments (scFvs, 25 kDa). Their diminutive size allows for the engineering of bivalent Fc fusion proteins that are half the size of a conventional antibody. While VNARs have only two CDRs, compared to six in humans and three in camelid domains, the CDRs protrude significantly, granting access to epitopes inaccessible to conventional antibodies. VNARs also possess two hypervariable loops in their framework regions, further increasing their binding capabilities and compensating for the loss of a CDR (32). VNARs are also easy to produce in large quantities from bacteria, are thermostable, and require little humanization due to their high homology with human light chains (33). These factors together have led to an increased interest in VNARs as targeting vectors for a variety of conditions (34–36).

Here, we developed a theranostic agent for MET-expressing NSCLC using a VNAR domain identified from the direct immunization of a nurse shark. We identify a high-affinity VNAR, termed vMET1. As a bivalent Fc fusion protein, vMET1-Fc binds recombinant human MET and MET-altered NSCLC cell lines, and is quickly and specifically internalized into MET-expressing cells. *In vivo* experiments confirmed ability of our VNAR to image both METex14-mutated and MET-amplified xenograft models of lung cancer by positron emission tomography/computed tomography (PET/CT) and single-photon emission computed tomography (SPECT). An anti-cancer effect was observed in our xenograft models treated with [^177^Lu]Lu-vMET1-Fc, highlighting its potential success as a theranostic agent. Finally, a non-human primate (NHP) study demonstrated the safety of vMET1-Fc, showing it clears predictably and caused no major health issues as described by observation and blood analysis. Our findings further underscore the potential of VNARs as targeting vectors for the delivery of therapeutic payloads and serve as a milestone for an emerging theragnostic approach.

## Materials and Methods

All animal studies were conducted under protocols approved by the University of Wisconsin Institutional Animal Care and Use Committee: M006482-R01-A01 (mouse), M006481-R01 (shark), and G0006671 (NHP). Macaques were provided by the Wisconsin National Primate Center. All cell experiments were performed within 4 months of thawing cell lines from frozen cell stocks. Cell lines were authenticated using short-tandem repeat analysis and routinely monitored for mycoplasma contamination prior to our studies.

### Study design

This study was to evaluate the potential of a MET-targeting, VNAR-based antibody as both a diagnostic and therapeutic tool. Studies begin with the initial generation and discovery of vMET1, follow its *in vitro* and later *in vivo* characterization in both mice and non-human primate (NHP) models. *In vitro* experiments were performed at least twice independently, and all cell-based imaging studies used at least three technical replicates within independent runs. *In vivo* studies were performed once. The sample size for *in vivo* mouse experiments was decided using our previous experience with this method, as well as an effort to use the smallest number of animals necessary. Mice were randomly placed in study groups at the start of each study, but researchers were not blinded during subsequent data collection. NHP studies were performed with *n* = 2 per group due to limited primate availability.

### Next-generation sequencing of the MET-immunized phage display library

Next-generation sequencing (NGS) analysis was performed on VNAR clones from the immunized library to further validate the immunization and clarify the composition and degree of diversity in the library. To determine the fraction of the library with sequences unique to this immunization, a sequence alignment analysis was performed using sequence data from the library in parallel with NGS data from an immunized library from an unrelated antigen as a control library. The control library had a similar number of full-length VNAR reads as the MET-immunized library, with <0.4% of sequences being shared by both libraries. VNARs were typed (I-IV) by established methods (37) using the number and arrangement of non-canonical cysteine residues, and the presence or absence of a conserved tryptophan residue adjacent to a CDR1 cysteine.

### ELISA

VNARs were produced for 192 clones using a 5 mM IPTG induction in microtiter plates. Soluble hemagglutinin (HA)-tagged VNARS which leaked into the culture media were screened for binding to MET by ELISA. MaxiSorp plates (Nunc Cell Culture, Thermo Fisher Scientific, Waltham, MA) were coated with 50 µL of 5 µg/mL streptavidin (Promega, Madison, WI) in PBS, overnight at 4° C. Wells were washed two times with PBS and blocked in 375 µL of 2% BSA in PBS for one hour, shaking, at room temperature. Wells were then washed three times with 0.005% Tween 20 in PBS, and all proceeding washes were done with this solution as well. Wells were then coated with 50 µL of biotinylated MET (1 µg/mL in PBS, 1% BSA, and 0.005% Tween 20). Plates were shaken at room temperature for one hour, then washed three times. Next, the supernatants of VNAR-induced cultures were added to each well and shaken at room temperature for 1 hour. Wells were washed three times, and then VNAR binding was detected with a 1:1000 dilution of anti-HA-Peroxidase monoclonal antibody (Roche, Mannheim, Germany, 3F10) in PBS with 1% BSA. Finally, 50 µL 1-Step™ Ultra TMB-ELISA Substrate Solution (Pierce, Thermo Fisher Scientific, Waltham, MA) was added to each well. Reaction was stopped with 10 µL of 1 M H_2_SO_4_, and absorbance was read on a microplate reader (Tecan) at 450 nm. Clones with high absorbance signals were identified as MET binders and sent for Sanger sequencing. Dilution ELISAs were performed by 2-fold serial dilutions from IPTG-induced supernatant in order to confirm saturable binding. VNAR binding was detected as described above.

### Biolayer interferometry

All biolayer interferometry (BLI) assays were performed using an Octet R8 (Sartorius, Göttingen, Germany), in a 1% BSA (w/v) in PBS buffer. Mouse MET (MET-M52H3, Acro Biosystems) and macaque MET (MET-C52H9, Acro Biosystems), which were not purchased biotinylated, were each biotinylated using EZ-Link NHS-PEG_4_-Biotin (A39259, Thermo Fisher Scientific, Waltham, MA), according to vendor recommendations. *Screening shark plasma samples.* Biotinylated human MET was diluted to 50 nM and immobilized on Octet streptavidin SAX biosensors (Sartorius, Göttingen, Germany) for 60 seconds. Sensors were then exposed to plasma samples from each blood draw timepoint, diluted 1:200 in assay buffer. IgNARs were allowed to associate with MET for 10 minutes, and then dissociated in assay buffer for 10 minutes. *Determination of purified antibody dissociation constants.* Pre-hydrated SAX biosensors were equilibrated for 60 seconds in assay buffer, then loaded with 50 nM MET (human, mouse, or macaque), followed by another 60-second baseline in assay buffer. Sensors were then exposed to serially diluted vMET1 monomer or vMET1-Fc, and then to pure assay buffer to measure dissociation. The exact length of loading, association, and dissociation steps was optimized to each protein and assay. Each experiment included two control wells: one in which unloaded SAX sensors (i.e. no MET) were exposed to analyte protein at the highest concentration used in the assay, to confirm a lack of binding to empty sensors. Second, a sensor was loaded with MET but exposed only to buffer, with no analyte included, to ensure the specificity of signal on MET-loaded sensors. Binding affinities were determined in the Octet Data Analysis software (v12.0.2.3), using conditions for a bivalent analyte (excepting the monomer assay) and background subtracting data from reference wells. The vMET1 monomer quickly reached a steady state at reasonable concentrations, and so a steady-state dissociation constant could be used. The bivalent Fc required a kinetic analysis using association and dissociation rates, as it would not reach a steady state in a reasonable assay. *Antibody epitope binning with HGF.* Hydrated SAX biosensors were subjected to a 60-second baseline in assay buffer, and then loaded with biotinylated human MET (50 nM, 100 s), followed by the establishment of a second baseline. Then, sensors were loaded with a saturating concentration of vMET1-Fc until a stable signal was achieved (1 µM, 180 s). This was followed by an extended exposure to recombinant human Hepatocyte Growth Factor (Peprotech, Thermo Fisher Scientific, Waltham, MA) for 30 minutes, at 50 nM. The HGF well also had 1 µM vMET1-Fc to ensure the sensor remained saturated. Matching dissociations followed the association for equal lengths of time. This assay was compared to one in which HGF was associated with MET in the absence of vMET1-Fc.

### RT-qPCR

Total RNA was isolated from cultured cells using TRIzol reagent and was subsequently treated with DNase1 to remove genomic DNA contamination. RNA concentration and quality were assessed by NanoDrop 2000 spectrophotometer. First-strand cDNA synthesis was performed using iScript cDNA Synthesis Kit (BioRad) with 1 ug RNA. Quantitative real-time PCR (qPCR) was carried out on QunatStudio3 (Applied Biosystems) using SYBR Green Dye master mix (PowerUp, Thermo Fisher). Primer pairs were purchased from IDT: MET (forward primer: 5’-CAGCCATAGGACCGTATTTCGG-3’ and reverse primer: 5’-TGCACAGTTGGTCCTGCCATGA-3’) and GAPDH (forward primer: 5’-GTCTCCTCTGACTTCAACAGC and reverse primer: 5’-ACCACCCTGTTGCTGTAGCCAA-3’). Each transcript was analyzed in triplicates. GAPDH was used as reference gene for normalization. PCR reactions (10 μL) were set up for amplification and thermal cycling conditions included an initial denaturation at 95 °C for n, followed by 40 cycles of 15 sec at 95 °C for denaturation and 1 min at 60 °C for annealing/extension, and melt curve. The results by the ΔΔCt method were analyzed by Design and Analysis 2 software (QuantStudio).

### MET copy number variation assay by qPCR

Genomic DNA was isolated from cultured cells using Wizard gDNA isolation Kit (Promega) according to the manufacturer’s instructions with RNase A treatment to remove RNA contamination. DNA was quantified by NanoDrop 2000 spectrophotometer. MET copy number variation was determined by predesigned TaqMan Copy Number Assay (Life Technologies, Hs04993403_cn; FAM-dye labeled) and TaqMan Copy Number Reference Assay as RNaseP (Life Technologies, #4403328, VIC-dye labeled). qPCR was performed using QuantStudio3 Real-Time PCR System (Applied Biosystems). Samples were normalized to a reference gene, RNAseP, known to have two copies, then compared to the calibrator, a normal human genomic DNA with a known, stable copy number of two (TaqMan Control genomic DNA, #4312260, Life Technologies). The TaqMan PCR reaction mixture was assembled in a final volume of 10 μL with TaqMan Genotyping Master mix, TaqMan MET Copy Number Assay or TaqMan Copy Number Reference Assay (RNaseP), and 10 ng of the genomic DNA as the template. Samples were thermally cycled under the following conditions: 95°C for 10 min (one cycle); 40 PCR cycles of 95°C for 15 seconds and 60°C for 1 minute. All reactions were performed in quadruplicates. The PCR data using the ΔΔCt method were analyzed using CopyCaller software (Thermo Fisher) to determine the copy number variation.

### Receptor quantification via flow cytometry

Receptor density was quantified by flow cytometry using commercial Onartuzumab (A2468, Selleckchem, Houston, TX) targeting the MET receptor. The antibody was conjugated with Alexa Fluor 647 (AF647) using a protein labeling kit (Thermo Fisher, #88068), and the fluorochrome-to-protein (F/P) ratio was determined via spectrophotometry. To determine receptor density, cell line suspensions were prepared and incubated with a saturating concentration (200 nM) of AF647-Onartuzumab for 30 min. For quantification, a standard curve was generated using calibration beads with known AF647 binding sites (Quantum Alexa Fluor 647 MESF kit; Bangs Labs). The Mean Fluorescence Intensity (MFI) of the stained cells was converted to Molecules of Equivalent Soluble Fluorochrome (MESF) using the standard curve; this value was subsequently divided by the F/P ratio to calculate the absolute number of receptors per cell.

### Cell proliferation assay

Cells were plated in 96-well plates at growth rate optimized densities (3,000–10,000 cells/well). After overnight incubation, cells were treated with the indicated doses of vMET1-Fc or vehicle control (PBS). Following 72 hours of treatment, Cell Counting Kit-8 (Dojindo Molecular Technologies) was added to each well, and absorbance was measured at 450 nm using a SpectraMax i3 plate reader (Molecular Devices). Absorbance from the treated wells was normalized to the control wells, and half-maximal inhibitory concentration (IC50) values were calculated with GraphPad Prism (version 10.5.3).

### Immunofluorescent fixed-cell staining

Eight-well chambers (coverslip slide #1.5 thickness) were used to seed 40,000 to 80,000 cells per well depending on cell type, followed by an incubation at 37 °C overnight. Cells were incubated with 10 nM of Alexa647-labeled vMET1-Fc for 24 h, 4 h, or 5 min and pHrodo Green Dextran 10,000 MW (Thermo Fisher Scientific, P35368) for 2 h. After incubation, cells were washed with 1X PBS and were fixed with 4% paraformaldehyde for approximately 12 min. After fixation, cells were washed again three times with 1X PBS and stained with Hoechst 3342 at ratio 1:2000 PBS and CellBrite® Cytoplasmic Membrane Dye at ratio 1:200 PBS (Biotium, #30022) for 20 min. Cells were washed three times again with 1X PBS, coverslip was mounted with PBS and sealed with clear nail polish, and was immediately imaged with a Nikon Spinning Disk Confocal microscope at 60X with oil and Blue/Green/Red/Far Red filter settings. Quantification of vMET1-Fc signal overlap with endosome stain per cell was calculated by defining cell areas by doubling size of nucleus, and percentage of voxels positive for vMET1-Fc staining that were also positive for endosome staining (threshold fluorescent for both channels 0.3).

### Internalization of pHrodo red labeled antibody constructs

Cells were seeded 3000 to 10,000 cells per well depending on cell type. vMET1-Fc was labeled using pHrodo Red (Thermo Fisher Scientific P36600) succinimidyl ester prior to Incucyte imaging. Cells were treated with indicated concentrations ranging from 0.01 nM to 250 nM with their respective cell media treated with approximately 25 mM of HEPES. Plates were imaged using an Incucyte SX5 Live-Cell Analysis System (Sartorius, Göttingen, Germany). Images were taken every 4 hours for 72 hours total, using phase and orange imaging channels, a 400 ms acquisition time, and a 10x objective. Signal was measured as an integrated intensity above a set background threshold across five images per well, normalized to the cellular confluency of each image. During imaging, cells were maintained at 37° C and 5% CO_2_. Acquired images were analyzed with the Incucyte’s Analysis Wizard. Images were chosen for analysis at 12-hour intervals and analyzed with artificial intelligence confluence segmentation in the phase imaging channel. In the orange imaging channel, surface fit segmentation and a threshold of 0.6 OCU (after background subtraction) was used, with edge split off. After analysis, images over time were plotted as integrated orange intensity per image, normalized by division to confluence area of the image (OCU x µm^2^/Image / %), along with standard error per well at each timepoint.

### PET/CT imaging

Mice bearing with EBC-1, UW-Lung-21, and T-47D tumors were injected with 5.55 MBq of [^89^Zr]Zr-vMET1-Fc by tail vein injection. Mice (n=4/ cell line) were anesthetized by inhaling 2.5% isoflurane. Images were acquired at 4, 24, 48, 72, and 96 h timepoints by an Inveon micro-PET/CT Scanner (Siemens Medical Solutions). 80 million counts using a γ-ray energy window of 350−650 keV and coincidence timing window of 3.428 ns were acquired for PET list mode data. A CT-based attenuation correction was performed for approximately 10 min with 80 kVp, 1 mA, 220 rotation degrees in 120 rotation steps, 250 ms exposure time and subsequently reconstructed using a Shepp-Logan filter with 210-μm isotropic voxels. Scans were reconstructed using three-dimensional ordered-subset expectation maximization (2 iterations, 16 subsets) with a maximum *a posteriori* probability algorithm (OSEM3DMAP). Two-dimensional (2D) images and maximum intensity projections (MIPs) were analyzed, and the region of interest (ROI) was manually drawn using the Inveon Research Workspace (Siemens Medical Solutions). ROI analysis of the images was used to determine tracer uptake in tumor and heart, and quantitative results are given as a percentage injected dose per gram of tissue (%ID/g). An *ex vivo* biodistribution measurement was carried out after the last timepoint. Major organs, tissues, and blood were collected, and the radioactivity in each sample was measured by using a Hidex automatic gamma counter (Hidex Oy, Turku, Finland). Count data were background- and decay-corrected, and %ID/g for each tissue sample was calculated by normalization to the total amount of activity injected into each mouse.

### SPECT/CT imaging

Mice bearing subcutaneous EBC-1 and UW-Lung-21 tumors were administered 11.1 MBq of [^177^Lu]Lu-vMET1-Fc by tail vein injection. Longitudinal SPECT/CT imaging was performed using a MILabs U-SPECT6/CTUhr system (Houten, The Netherlands). Mice (n = 4/cell line) were anesthetized with 2% isoflurane in oxygen and positioned prone for imaging sessions conducted at 4, 24, 48, 72, and 120 h post-injection. CT scans were acquired for anatomical reference and attenuation correction, and subsequently fused with SPECT images. SPECT data were reconstructed using a similarity-regulated ordered-subset expectation maximization (SROSEM) algorithm. Quantitative image analysis was performed by manually delineating volumes of interest (VOIs) on the fused CT/SPECT images over the tumor and heart. The radiotracer uptake in each tissue was calculated and expressed as the percentage of injected dose per gram of tissue (%ID/g). An *ex vivo* biodistribution measurement was carried out after the last timepoint. Major organs, tissues, and blood were collected, and the radioactivity in each sample was measured by using a Hidex automatic gamma counter (Hidex Oy, Turku, Finland). Count data were background- and decay-corrected, and %ID/g for each tissue sample was calculated by normalization to the total amount of activity injected into each mouse.

### Xenograft growth delay studies

Mice were injected with tumor cells (EBC-1 or UW-Lung-21) mixed in Matrigel subcutaneously above the right shoulder blade. When tumors grew to a mean volume of 150 to 300 mm^3^, mice were randomized into groups and treated with either control (isotype), vMET1-Fc, or 11.1 MBq [^177^Lu]Lu-vMET1-Fc. A single dose of each treatment (1 mg/kg) was injected by tail vein. Tumor volume was measured twice weekly with digital vernier calipers and calculated using *V* = (π/6) × (large diameter) × (small diameter)^2^. Each experimental group contained 8 to 10 mice.

### Acute toxicity animal study

Immunocompetent non-tumor bearing mice (ICR, ∼30 g, Jackson Laboratory) were randomized into groups and treated with control (isotype), vMET1-Fc, or 11.1 MBq [^177^Lu]Lu-vMET1-Fc. A single dose of each treatment (1 mg/kg) was injected by tail vein. Complete blood count (CBC), blood chemistry profiles, and histology evaluation were performed at 3, 7, and 30 days post-injection. Blood was collected at each timepoint by cardiac puncture as a terminal procedure. CBC analysis was performed on 50 µL EDTA-treated blood using a Vetscan HM5 analyzer (Zoetis Diagnostics). Blood chemistry analysis was conducted with 100 µL plasma separated from lithium heparin-treated blood (4000 x g, 10 min) and evaluated using a Vetscan VS2 analyzer with the Preventive Profile Care Plus rotor, encompassing 15 parameters (Zoetis Diagnostics). Additionally, liver, kidney, and spleen tissues were harvested at each timepoint for histological examination using H&E staining. Histopathologic review was performed by a board-certified veterinary pathologist on glass slides. Then, slides were scanned by Aperio Digital Pathology Slide Scanner, AT2 (Leica Biosystems) at 40X. Image analysis for the spleen was conducted using HALO AI (v3.6.4134.226; Indica Labs). HALO AI Preset Tissue Detection and QC Slide V2 were applied to all slides to identify normal spleen and exclude any artifacts (e.g., folds, crushing). Areas with additional artifacts not identified by QC Slide V2 were manually excluded. For classifier validation, 28 specimens (representing all treatment groups) were selected. A pathologist annotated 5X fields (>7 mm2) to define tissue classes: red pulp, white pulp, and background. A DenseNet V2 convolutional neural network (CNN) was trained at 2 µm/pixel resolution and a minimum object size of 200 um2 for >40,000 iterations, achieving a cross-entropy 0.211 and a total F-1 score of 0.96205. The validated classifier was applied to all WSI. The white pulp and red pulp areas were quantified across one longitudinal section to calculate the white pulp-to-red pulp ratio.

### Mouse immunogenicity

Eight to ten-week-old immunocompetent C57BL/6 mice (n=4 mice/sex/experimental group) were injected subcutaneously with 1 mg/kg of antibody weekly for 7 weeks with vMET1-Fc, commercial MET antibody onartuzumab (A2468, Selleckchem, Houston, TX), or PBS. One week after the final dose, mice were sacrificed and blood was collected via axillary bleed. Blood was centrifuged to separate its components. Plasma was aliquoted and flash frozen, while red blood cells were discarded. Clear MaxiSorp plates (Nunc Cell Culture, Thermo Fisher Scientific, Waltham, MA) were coated with 50 µL of 5 µg/mL streptavidin in PBS. Plates were incubated overnight at 4 °C. Plates were then washed three times with 0.1% Tween 20 in PBS (PBST) before adding 4% (w/v) skim milk powder in PBS (M-PBS) and blocking for 1 hour, shaking at room temperature. Three PBST washes followed this step and all subsequent steps.

Biotinylated vMET1-Fc or onartuzumab were added to each well, at 1 µg/mL in 2% M-PBS, and incubated for 1 hour. Meanwhile, plasma samples were serially diluted (1:4) in 300 mM acetic acid and left at room temperature for 1 hour to dissociate any immunocomplexes. Before sample application, wells were neutralized with 1 M Tris, pH 9.0. The acid-treated plasma samples were applied to Maxisorp plates and incubated for 2.5 hours to allow binding. A 1:10,000 dilution of anti-mouse IgG (H+L)-peroxidase antibody produced in goat (Thermo Fisher Scientific) was made in 4% M-PBST and incubated for 1 hour. Colorimetric signal development was induced using 1-Step™Ultra TMB-ELISA substrate (Thermo Fisher Scientific) and neutralized with 1 M sulfuric acid. Colorimetric intensity was read using an Infinite M200 pro plate reader (Tecan, Switzerland) at 450 nm.

### Pharmacokinetics of [^89^Zr]Zr-vMET1-Fc in NHP

Two healthy male macaques (*Macaca fascicularis*) aged 5.5 and 6 years, and weighing 10.25 and 8.95 kg, respectively, were selected for this study. The pharmacokinetics and distribution of [^89^Zr]Zr-vMET1-Fc was evaluated in a PET/CT study. Animals were administered, respectively, 104.7 MBq and 105.8 MBq of [^89^Zr]Zr-vMET1-Fc (0.2mg/kg) as an intravenous bolus. Subsequently, imaging was performed at 1, 24, 72, and 144 hours post-injection. ROI analysis was performed manually using the Inveon Research Workspace. ROI analysis of the images was used to determine tracer uptake in major organs or tissues, and quantitative results are given as standardized uptake values normalized to body weight (SUV_bw_).

### Analysis of NHP blood samples

One week after initial injection with [^89^Zr]Zr-vMET1-Fc, macaques were both dosed with cold vMET1-Fc at 1 mg/kg, weekly for three weeks, with blood draws performed concurrently. Blood samples were collected in both EDTA and Heparin (BD and Company, Franklin Lakes, NJ) coated collection tubes. A portion of blood samples from each draw within the first seven days was gamma counted using a Hidex Automatic Gamma Counter. Then, 1 mL of whole blood from the EDTA tube was used to perform a complete blood count (CBC) on a VETSCAN HM5 hematology analyzer. Resulting values were plotted over time against reference values provided by Zoetis via the VETSCAN HM5. After centrifuging the remaining blood, plasma from EDTA tubes was stored for later ELISA analysis, while plasma from heparin tubes was taken for immediate blood chemistry analysis. This plasma was loaded onto a VetScan® Preventative Care Profile Plus (Abaxis, #500-047) reagent rotor and measured on a VETSCAN VS2 blood chemistry analyzer. Resulting values were plotted over time against reference values provided by the Wisconsin National Primate Research Center based on the animals’ sex and weight class.

### Statistical analysis

Experiments conducted *in vitro* were repeated three times, and statistical analyses were carried out using a t-test or one-way ANOVA. Data was presented as the mean ± standard error of the mean (SEM). Statistical analysis of Progression-Free Survival (PFS) curves was performed with a log-rank test. A *P* value of < 0.05 was considered statistically significant. All graphs and *in vitro* analyses were performed and graphed using GraphPad Prism version 10.0.3 (Prism) or RStudio (2021.09.0).

### Data Availability

The data generated in this study are available within the article and its supplementary data files.

## Results

### Immunization of a nurse shark generates high-affinity anti-MET VNARs

A juvenile male nurse shark was immunized (**Fig. 1A**) with the extracellular domain of recombinant human MET (aa 25-932) (**Fig. 1B and C**). Evidence of an immune response was measured by screening the shark plasma for anti-MET shark antibodies via biolayer interferometry (BLI). No binding was detected in naïve plasma, while subsequent samples showed a steady, time-dependent increase in MET association throughout the immunization series (**Fig. 1D**). Plasma collected at week 18 failed to show a response to control biosensors lacking MET. The shark immune repertoire was captured by cloning VNAR sequences present in buffy coat isolated from blood draws. VNAR-encoding sequences were amplified and used to create a VNAR phage display library (1.9 x 10^8^ cfu), which was then screened against MET. Next-generation sequencing (NGS) analysis of the immunized library showed that the control library had a similar number of full-length VNAR reads as our MET-immunized library, with <0.4% of sequences being shared by both libraries (**Fig. 1E**). Nurse sharks are unique in that they possess all four VNAR types. Plotting the NGS results demonstrated that the library possessed diversity across the four different VNAR types and had varying CDR3 lengths (**Fig. 1F**).

**Figure 1.**
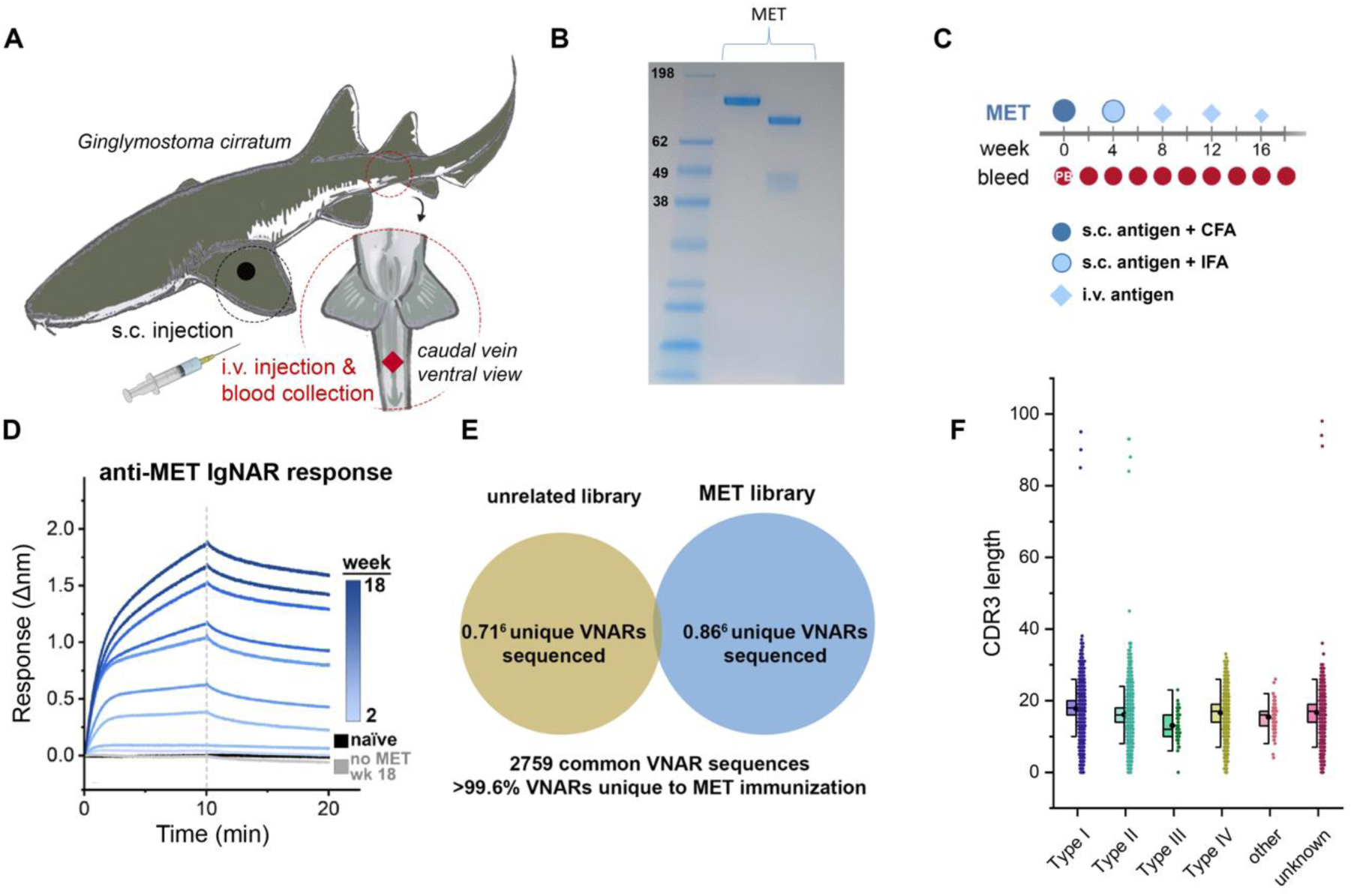
Nurse shark immunization and characterization of resulting phage-display library. **A)** Designation of sites used for subcutaneous (s.c.) and intravenous (i.v.) delivery of immunogens and blood collection. **B)** SDS-PAGE gel with Coomassie staining of extracellular domain of the MET protein with a See Blue ladder (left), the MET protein (center) and the MET protein under reducing conditions (right). **C)** Diagram of the time course, injections, adjuvants, and blood collections throughout the MET immunization program. **D)** Biolayer Interferometry (BLI) sensorgram from a representative experiment demonstrating the mobilization of an anti-MET immune response after MET immunization. Diluted plasma samples were collected over time and screened against biosensors loaded with MET to detect the presence of MET-binding antibodies. **E)** Venn diagram of sequence overlap between NGS datasets derived from sequencing the MET-immunized VNAR phagemid library and a library constructed for an unrelated immunogen. **F)** A column scatter of the number of amino acids in the CDR3 of VNARs by subtype, overlaid with box plots to illustrate the spread of the data.

### Identification of anti-MET VNARs

After two rounds of phage display biopanning, 192 clones were screened using a monoclonal ELISA assay. Clones that showed strong binding in this assay were selected for further analysis. The eleven most promising clones were tested for concentration dependent, saturable binding using a serial dilution ELISA (**Fig. 2A**). Sanger sequencing was used to verify legitimate VNAR sequences and eliminate redundant clones. Based on sequence analysis and preliminary expression yields, a VNAR termed vMET1 was selected for further analysis. After investigating the sequence of the VNAR and generating an initial structural prediction (Supplementary Fig. S1A and B), we found that vMET1 closely resembled a type II VNAR, but has an extra non-canonical cysteine in its HV2 region (Supplementary Fig. S1C). vMET1 was expressed in bacteria as a monomer, purified (Supplementary Fig. S2A), and analyzed by BLI to determine the affinity (**Fig. 2B**). By BLI, we next determined that the vMET1 monomer had a K_D_ of 34 nM, for human MET; however, it also had an unfavorable fast dissociation rate. To create a construct with high affinity and a slow dissociation rate, vMET1 was expressed as human IgG1 fragment crystallizable (Fc) fusion protein (Supplementary Fig. S2B and C). The resulting bivalent vMET1-Fc had a dramatically improved dissociation rate and a K_D_ of 26 pM (**Fig. 2C**). We next determined the cross-reactivity of vMET1-Fc for murine and cynomolgus monkey MET. While vMET1-Fc showed no binding to murine MET (Supplementary Fig. S3A), it did exhibit strong binding to macaque MET, with a K_D_ of 372 pM (Supplementary Fig. S3B). Finally, the specificity of vMET1-Fc was assessed by screening for binding against a panel of cell surface proteins, none of which bound vMET1-Fc (**Fig. 2D**).

**Figure 2.**
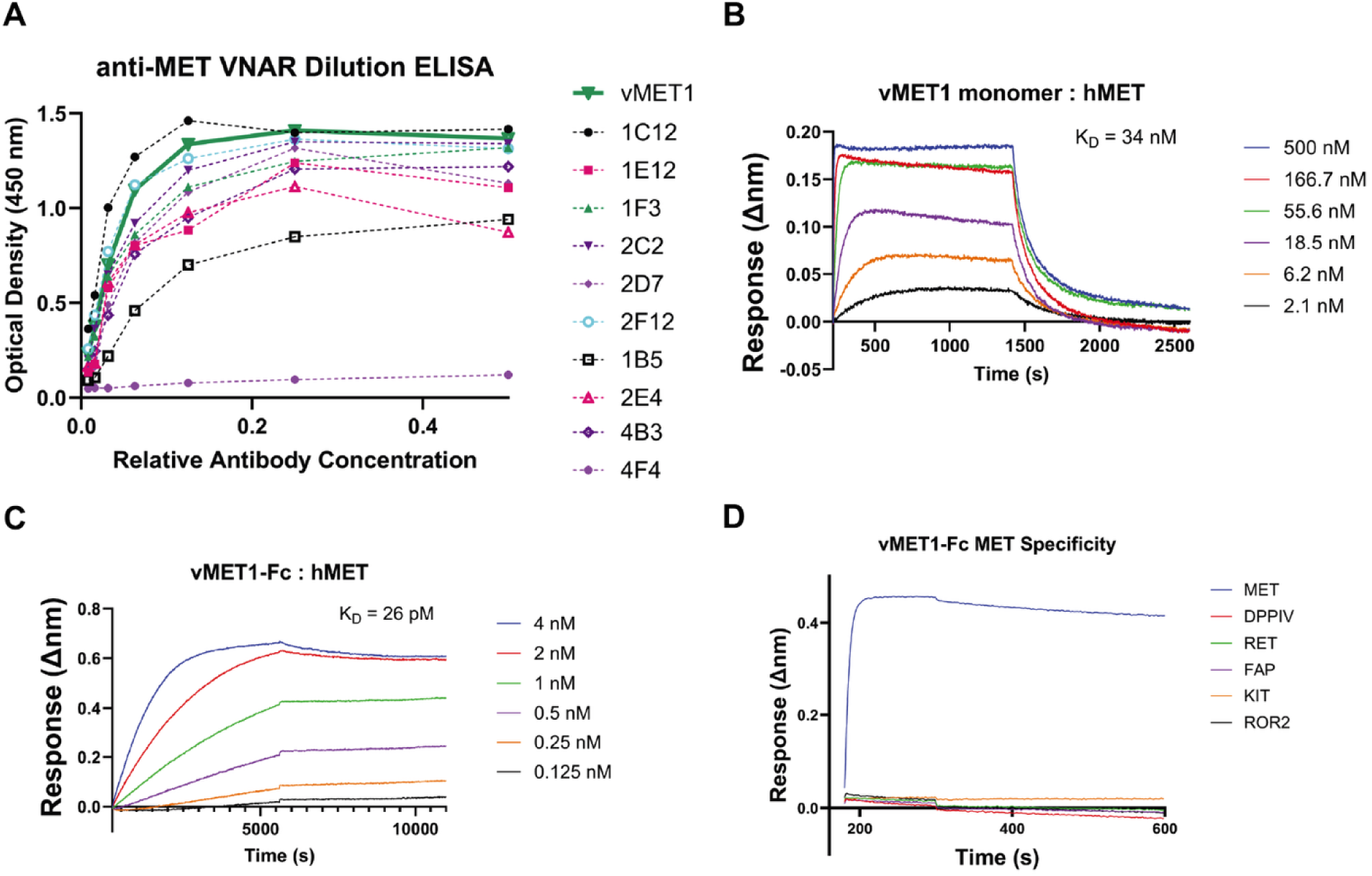
Discovery and assessment of potential anti-MET VNARs from a phage-display library. **A)** Dilution ELISA of promising VNAR clones demonstrate saturable binding to MET. Eleven unpurified VNARs were serially diluted and added to plates coated with MET protein. **B)** BLI sensorgram of sensors loaded with a variety of proteins, each tested against a standard concentration of vMET1-Fc. **C)** BLI sensorgrams of sensors loaded with human MET exposed to serially diluted vMET1 monomer or **D)** vMET1-Fc, followed by dissociation in assay buffer. Dissociation constants (K_D_) are listed for each assay.

### vMET1-Fc has high sensitivity and specificity for MET-expressing cells

A library of cell lines was curated and characterized based on origin site and co-occurring oncogene mutations. MET and phosphorylated-MET (p-MET) protein expression were evaluated via western blotting (**Supplementary Fig. S4A**). Next, the binding of vMET1-Fc to MET was measured in each cell line using flow cytometry, and a fold change in binding was calculated by comparison to the T-47D, a MET-negative breast cancer cell line (**Supplementary Fig. S4B**). MET expression characteristics of each cell line were assessed via quantification of MET RNA expression, MET gene copy number (CN), total MET and p-MET protein abundance, and MET receptor density (**Fig. 3A**). A strong positive correlation was observed between vMET1-Fc binding and all MET expression characteristics (**Fig. 3B**). The strongest associations were found with both MET receptor density (R = 0.82) and p-MET expression (R = 0.82). Strong positive correlations were also observed with MET RNA expression (R = 0.74) and total MET protein expression (R = 0.62), and a moderate positive correlation was found with MET CN (R = 0.58). From this panel, NSCLC MET-amplified cell line EBC-1 (38) and METex14-mutated cell line UW-Lung-21 (39, 40) were selected for further *in vitro* and *in vivo* analysis.

**Figure 3.**
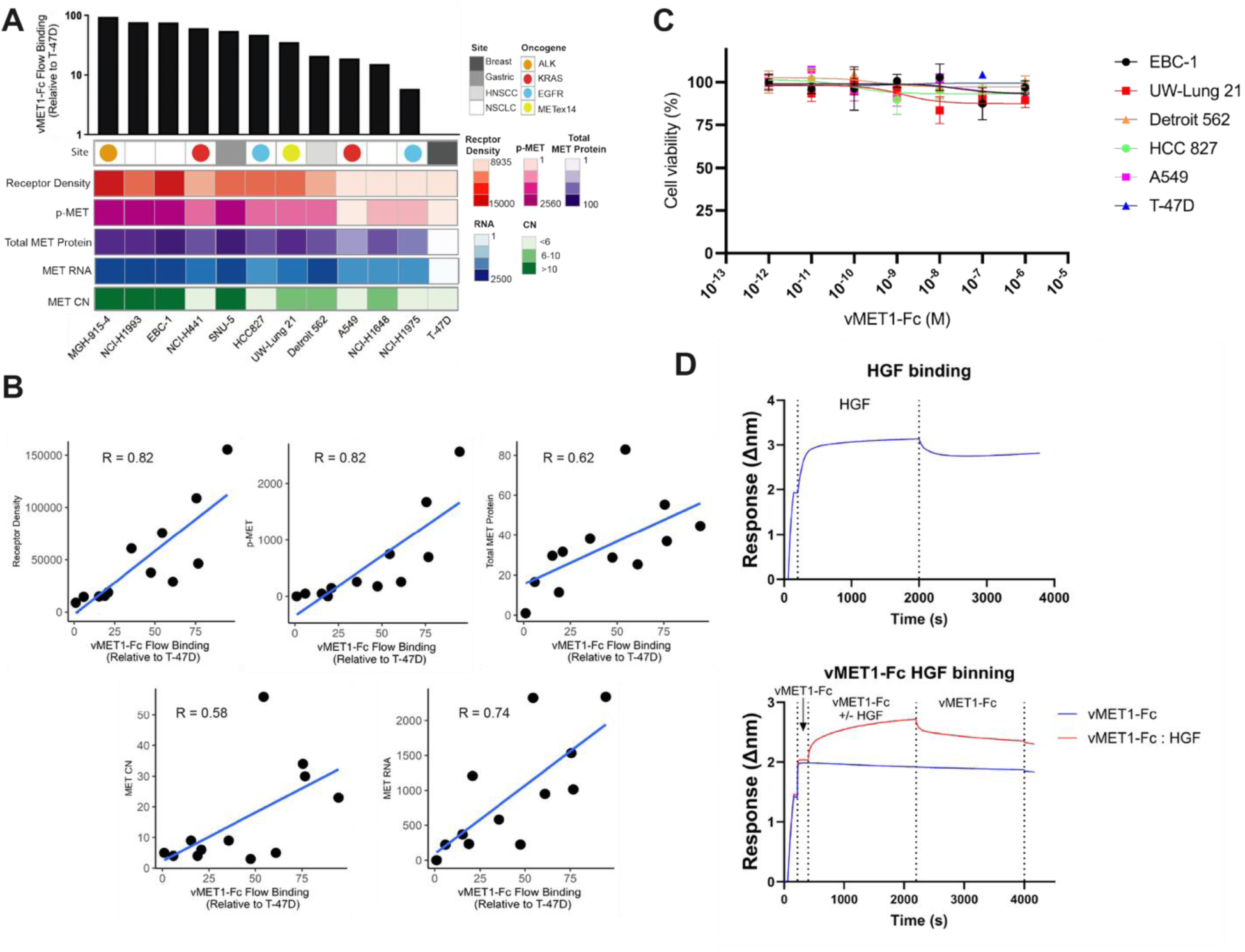
vMET1-Fc binding affinity is correlated with MET protein expression across cancer cell lines, and does not affect cell viability due to lack of competitive binding with HGF nor inhibition of MET downstream signaling. **A)** Heat map detailing in order top to bottom: site of origin and presence of oncogene, phosphorylated MET (p-MET) with respect to T-47D, MET Copy Number (CN), total MET protein expression with respect to T-47D, MET RNA, and MET receptor density. Numerical color spectrum is log-2 scaled. Black bar graph on top demonstrates vMET1-Fc binding quantified via flow cytometry normalized to T-47D. **B)** Individual scattered dot plots and linear regression modelling of vMET1-Fc binding with respect to receptor density, p-MET, total MET protein, MET RNA, and MET CN. Pearson correlation coefficient (R) was calculated for each variable comparison. Strong correlation was observed between antibody binding and MET receptor density. **C)** Viability assay to assess proliferation with concentrations of vMET1-Fc ranging from 1 fM to 1 µM does not show impact on cell survival across indicated cell lines. Points mean; bar SEM (n = 6). **D)** Sensorgram showing the binding of HGF to sensors loaded with human MET protein, and then dissociating in assay buffer (top) and the same assay, but vMET1-Fc is allowed to associate with sensors before the addition of HGF (bottom).

### vMET1-Fc does not affect cell viability nor downstream MET signaling

Our panel of MET-targeting antibodies was screened for their ability to modulate cell proliferation. vMET1-Fc, up to 1 µM, had no effect on cell viability across all assessed cell lines (**Fig. 3C**). To evaluate the impact of vMET1 on MET’s interaction with its sole known ligand, hepatocyte growth factor (HGF), we used BLI to compare HGF binding in the presence or absence of vMET1-Fc. No appreciable difference in HGF binding was observed (**Fig. 3D**), consistent with the expectation that HGF, a large protein, engages MET over a substantially larger surface area (41) compared to our VNAR, vMET1. Using the small-molecule MET tyrosine kinase inhibitor capmatinib as a positive control, western blot analysis confirmed that vMET1-Fc neither inhibited MET nor affected downstream signaling in MET-amplified EBC-1 cells, METex14 UW-Lung-21 cells, or MET-negative T-47D cells (Supplementary Fig. S4C).

### vMET1-Fc internalizes into MET-expressing cells

To measure the ability of the construct to internalize, vMET1-Fc was labeled with the pH sensitive fluorophore pHrodo Red to monitor movement through the increasingly acidic endolysosomal system. vMET1-Fc readily internalized into MET-expressing NSCLC cell lines, reaching maximum integrated intensity between 48 and 72 hours, while MET-negative T-47D cell showed negligible fluorescence (**Fig. 4A**). Approximate internalization half-lives were 22.0 h for EBC-1 and 35.3 h for UW-Lung-21 cell lines. Internalization was also assessed in a MET wild type immortalized renal cell line, RPETC/TERT1. Limited uptake was observed in RPETC/TERT1 as the intensity of the signal was much lower than the MET-altered cancer cell lines and an estimated internalization half-life of 40.0 h was measured. A panel of MET-expressing cell lines treated with varying concentrations of pHordo Red labeled vMET1-Fc demonstrated that signal intensity correlated with MET expression, and nonspecific internalization can be observed at concentrations above 100 nM (Supplementary Fig. S5A and B). Additional immunofluorescent (IF) imaging confirmed the selective binding and internalization of vMET1-Fc in the MET-positive cell lines (**Fig. 4B**). Cell plasma membrane (red) and endosomal (green) stain visualize the binding of the vMET1-Fc to cell surface receptors upon immediate exposure with gradual internalization, overlapping with green endosomal signal over time. Quantification of percent vMET1-Fc-Alexa Fluor 647 signal overlap with green endosomal signal across cell lines and time points affirmed the results observed with pHrodo red. MET-altered NSCLC cell lines demonstrated a significant increase in overlapped signals in MET-expressing cell lines, with EBC-1 cells demonstrating the highest percent overlap at 4 hours post treatment (**Fig. 4C**). Overlap signal was only quantified 24 hours post-treatment in RPETC/TERT1 cell lines and no significant overlap signal was observed in T-47D at any time points.

**Figure 4.**
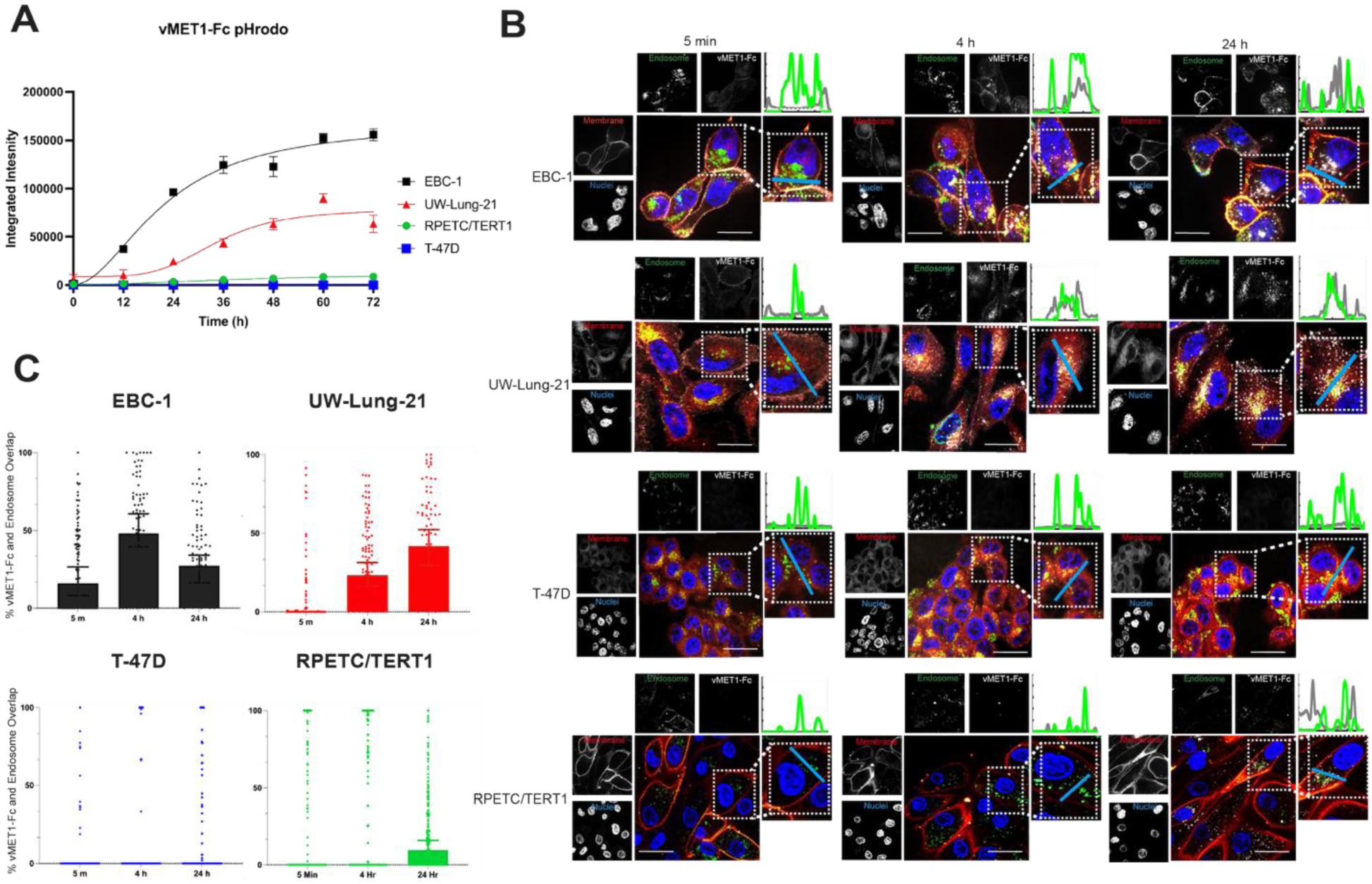
vMET1-Fc internalizes specifically into MET-expressing cell lines. **A)** Aggregate data from high-content live-cell imaging of vMET1-Fc internalization into MET-expressing EBC-1 and UW-Lung-21 cells and MET-negative T-47D cells. Antibody was directly labelled with pH-sensitive pHrodo Red, which increases fluorescence with decreasing pH, and resulting fluorescence was measured as integrated signal intensity normalized to confluency for three days. Points mean; bar SEM (n =5). **B)** Representative confocal microscopy images assessing vMET1-Fc internalization into MET-positive and -negative cell lines over time. Nuclei (blue), cell membranes (red), endosomes (green) and vMET1-Fc (white) are stained in large composite images, while separated channels are shown in smaller images. Blue line (inset) shows axis of the graph depicting vMET1-Fc and endosome signal at each time point. **C**) Quantification of percent overlap of vMET1-Fc color channel with endosomal color channel per cell (approximately 100 cells per time point per cell line) affirms internalization observed only within MET-expressing cell lines.

### [^89^Zr]Zr-vMET1-Fc as a PET/CT imaging agent

We next tested the ability of vMET1-Fc to detect MET expression using *in vivo* models of lung cancer by PET/CT imaging. The vMET1-Fc was radiolabeled with the positron-emitting radioisotope zirconium-89 (t_1/2_=3.7 d) using a deferoxamine (DFO) chelate that was site-specifically attached via click chemistry to two glycosylation motifs on the Fc domain. Mice bearing EBC-1, UW-Lung-21, or T-47D subcutaneous xenografts were injected with [^89^Zr]Zr-vMET1-Fc (7.4 MBq) and imaged serially out to 96 h post-injection. After analyzing 2D and maximum intensity projection (MIP) images of acquired scans, we found that vMET1-Fc showed significant and sustained uptake in MET-expressing tumors (**Fig. 5A**). Significant tumor uptake was evident by 4 hours post-injection in images with rapid clearance from the circulatory system and minimal normal tissue uptake in the MET-altered xenograft models. Region of Interest (ROI) analysis of heart and tumor over time quantified this observation, noting tumor-to-heart ratios of 4.0 and 4.6 in EBC-1 and UW-Lung-21 tumors respectively 96 h post-injection (**Fig. 5B**). A much lower tumor-to-heart ratio of 2.8 was observed in T-47D mouse models highlighting the sensitivity and specificity of [^89^Zr]Zr-vMET1-Fc *in vivo*. Additionally, *ex vivo* biodistribution analysis of multiple organs harvested at the 96 h timepoint show a much higher %ID/g in the tumor than any body organ or tissue, observing 20.0% ID/g and 30.2 %ID/g in EBC-1 and UW-Lung-21 tumors respectively. The next highest accumulation occurred as expected in clearance organs: the liver and kidneys (**Fig. 5C**).

**Figure 5.**
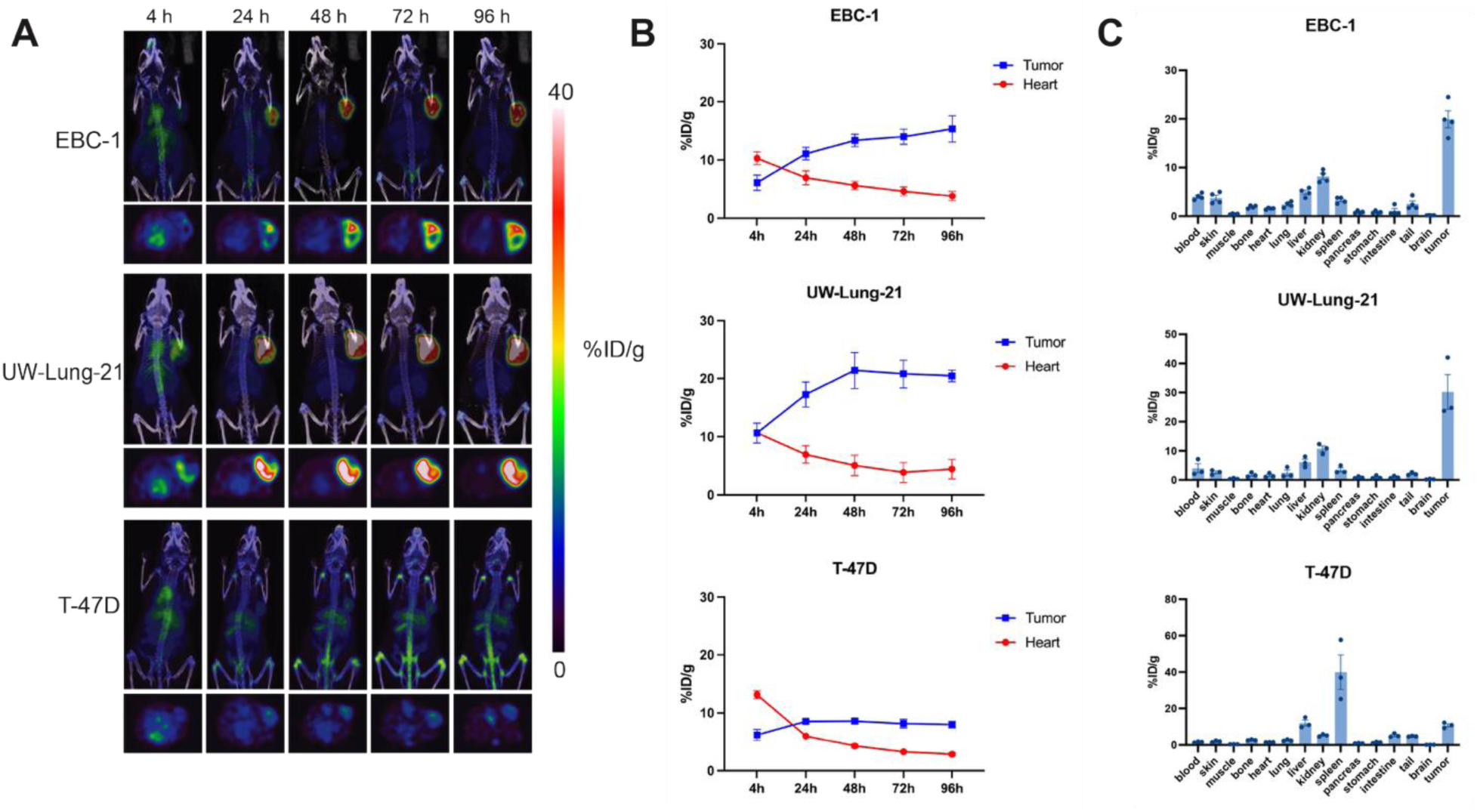
[^89^Zr]Zr-vMET1-Fc targets and accumulates in MET-expressing xenografts. **A)** Representative images from PET/CT scans of mice bearing xenografted tumors of EBC-1, UW-Lung-21, or T-47D cells (n = 4 per cell line) injected with [^89^Zr]Zr-vMET1-Fc and imaged at the given intervals. **B)** Region of interest (ROI) analysis of the PET images quantified radioactivity uptake (injected dose per gram (%ID/g)) across all time points for the heart and tumor. **C)** *Ex vivo* biodistribution analysis of [^89^Zr]Zr-vMET1-Fc activity across organs and tissue collected at 96 h post injection. There is significantly higher measured activity in MET-altered tumors compared to MET-negative tumors. Values are the mean (n = 4) ± SEM.

### [^177^Lu]Lu-vMET1-Fc as a radiotherapy

To assess the potential of vMET1-Fc as a targeted radiotherapy, the antibody was site-specifically radiolabeled with the beta particle-emitting radioisotope lutetium-177 (t_1/2_ = 6.7d) via a DOTA chelate group on the Fc domain. Approximately 11.1 MBq of [^177^Lu]Lu-vMET1-Fc were injected into mice bearing EBC-1 and UW-Lung-21 xenograft tumors. Single-photon emission computed tomography (SPECT) was performed starting at 4 h post-injection out to 120 h post-injection to detect the photon emitted by the decay of lutetium-177 (**Fig. 6A**). Imaging and ROI analysis of the tumors and heart demonstrated significant uptake in MET-altered lesions with rapid clearance from the circulatory system. The tumor-to-heart activity ratios were 4.4 and 3.5 in EBC-1 and UW-Lung-21 models respectively at 24 h post-injection (**Fig. 6B**). Biodistribution analysis at 120 h recapitulated the SPECT imaging data showing high tumor uptake and low retention in normal tissues (**Fig. 6C**). Dosimetry performed on Monte Carlo-supported software RAPID estimated tumor dose delivery of approximately 20.2 and 18.2 Gy for UW-Lung-21 and EBC-1 tumors respectively (fig. S6A). Imalytics was also used to affirm dose profiles to tumors as well as generate dose profiles for additional organs at risk (Supplementary Fig. S6B to D). The single dose of [^177^Lu]Lu-vMET1-Fc significantly delayed tumor growth and improved survival in both xenograft models compared to isotype control and cold vMET1-Fc (Supplementary Fig. S7A). The mean tumor doubling time (TDT) for EBC-1 tumors was 5.7 days for isotope-treated mice, and 6.2 days for vMET1-Fc-treated mice compared to 11.9 days in mice that received [^177^Lu]Lu-vMET1-Fc, (*P* < 0.0001 both comparisons, **Fig. 6D**). For UW-Lung-21 tumor xenografts, the mean TDT was 6.5 days in both the isotope-treated and vMET1-Fc treated mice, compared to 42.9 days in mice treated with[^177^Lu]Lu-vMET1-Fc, (*P* < 0.0001, both comparisons. **Fig. 6D**). No significant weight loss or sickness was observed in the treated mice (fig. 7B). Mice that received [^177^Lu]Lu-vMET1-Fc showed significantly improved overall survival compared controls (**Fig. 6E**). Specifically, the median survival time increased twofold in EBC-1 (47 days vs. 19 days) and three- to fourfold in UW-Lung-21 (37 days vs. 9–13 days), compared to mice that received vMET1-Fc (*P <* 0.001). Histological assessment of the liver and kidney revealed no evidence of damages caused by treatment of [^177^Lu]Lu-vMET1-Fc. However, validated AI analysis demonstrated a reduced white pulp-to-red pulp ratio in the spleens of mice treated with [^177^Lu]Lu-vMET1-Fc at day 30, correlating to histologically visual lymphodepletion of the white pulp and expansion of extramedullary hematopoiesis within the red pulp, which is expected with targeted ^177^Lu therapy (Supplementary Fig. S8). Complete blood count and blood chemistry was performed on mice harvested on days 3, 7, and 30. While transient changes in some metrics were noted among lutetium treated mice, all counts returned to levels comparable to controls within 30 days (Supplementary Fig. S9A and B).

**Figure 6.**
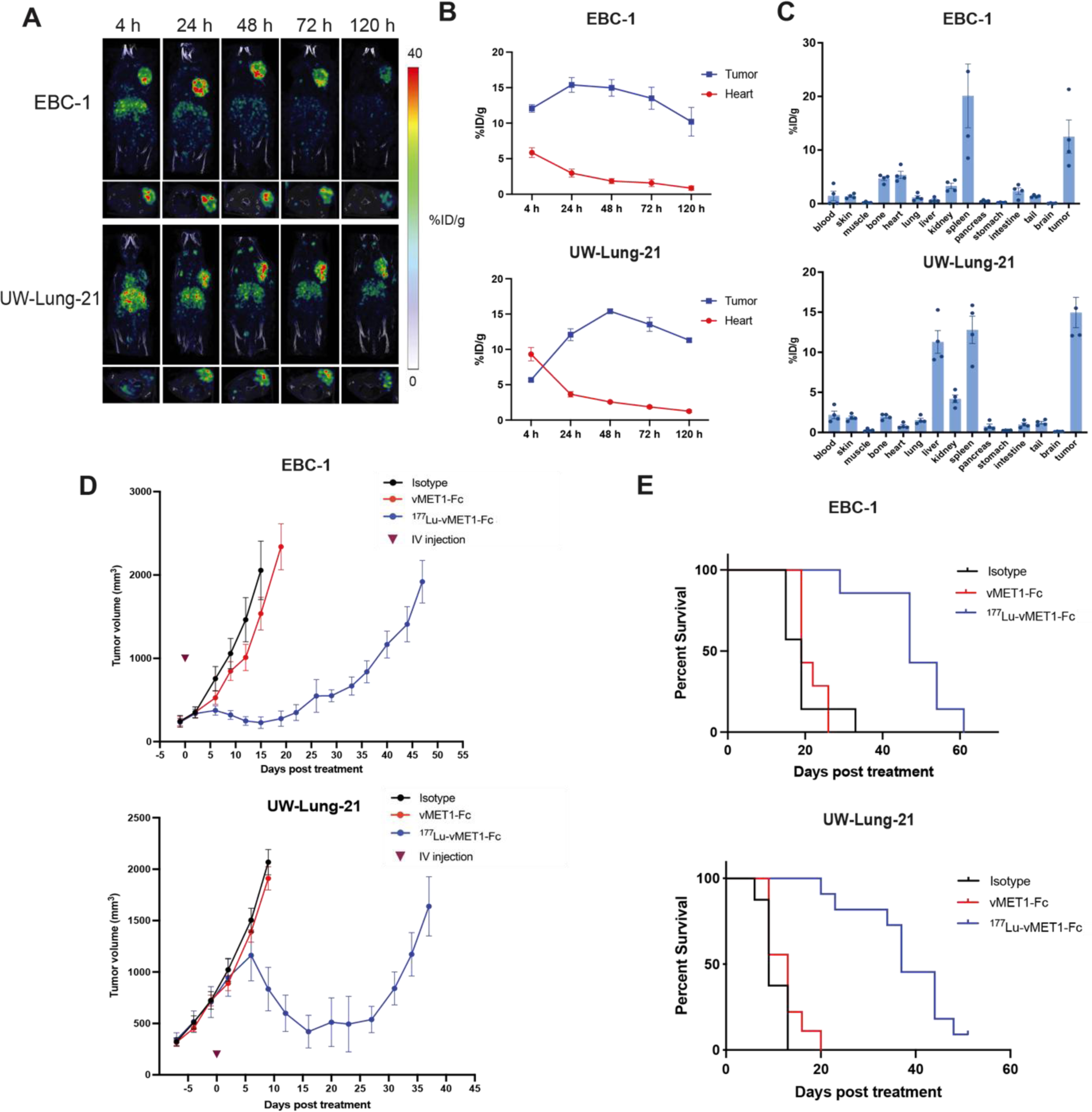
[^177^Lu]Lu-vMET1-Fc targets, accumulates and contributes to an anti-cancer effect in MET-expressing xenografts. **A)** Representative images from SPECT scans of mice bearing xenografted EBC-1 or UW-Lung-21 tumors (n=4) collected at the indicated time intervals after injection of [^177^Lu]Lu-vMET1-Fc. **B)** Region of interest (ROI) analysis of SPECT images quantified radioactivity uptake (injected dose per gram (%ID/g)) across all time points for the heart and tumor. Values are mean; bar SEM (n=4). **C**) *Ex vivo* biodistribution analysis of [^177^Lu]Lu-vMET1-Fc activity across organs and tissue collected 120 h post injection. Values are mean; dots are individual measurements; bar SEM (n=4). **D**) Tumor growth delay curves of EBC-1 or UW-Lung-21 xenografts treated with isotype (1 mg/kg), vMET1-Fc (1 mg/kg), or [^177^Lu]Lu-vMET1-Fc (11.1 MBq). A significant anti-cancer effect was seen when comparing [^177^Lu]Lu-vMET1-Fc to other groups. Mice were sacrificed when tumor volume exceeded 2000 mm^3^. **E**) Kaplan-Meier curve of mice from tumor growth delay experiment across groups. Survivability improved in mice treated with [^177^Lu]Lu-vMET1-Fc compared to other groups.

**Figure 7.**
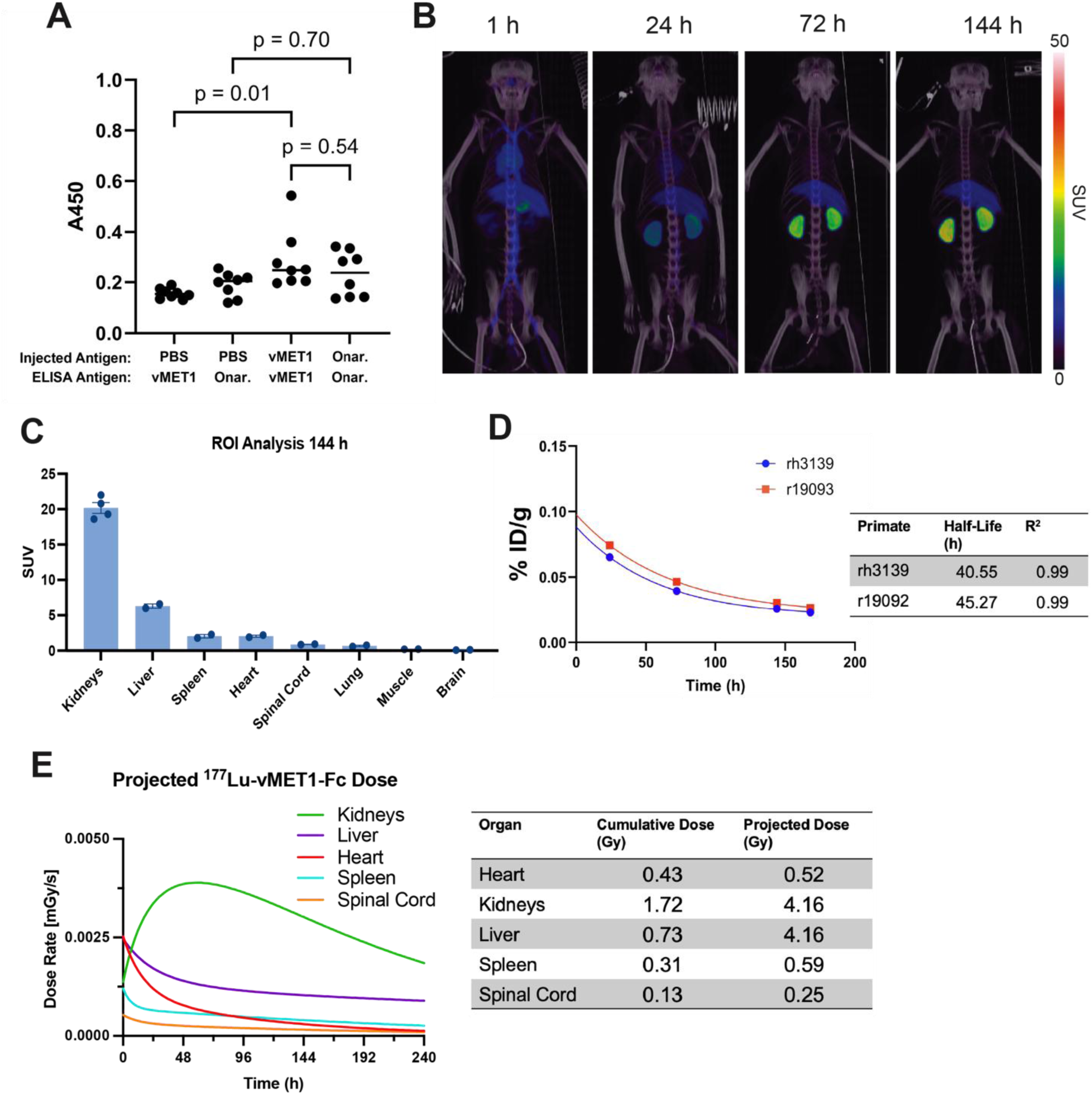
*In vivo* immunogenicity study to detect anti-drug antibodies in mice and macaques. **A**) Results of an ELISA run with diluted plasma of mice injected for seven weeks at 1 mg/kg with vMET1-Fc, commercial onartuzumab, or PBS. Left two groups detect anti-vMET1-Fc antibodies in mice injected with vMET1-Fc vs PBS, while the right two groups detect anti-onartuzumab antibodies in mice injected with onartuzumab vs PBS. **B)** Representative PET/CT scan of rhesus macaque (r19093) 1, 24, 72, and 144 h post-injection of [^89^Zr]Zr-vMET1-Fc. **C**) ROI analysis of identified organs at 144 h post-injection of [^89^Zr]Zr-vMET1-Fc. Values are mean; dots are individual measurements; bar SEM (n=2). **D**) Radioactivity measurement of blood samples collected at increasing timepoints post injection demonstrated rapid clearance of activity from circulatory system. Points are individual measurements; table estimates physiological [^89^Zr]Zr-vMET1-Fc half-life per primate and R^2^ fit of exponential decay function. **E**) Imalytics-derived dose rate profiles generated by applying the observed uptake of [^89^Zr]Zr-vMET1-Fc to a ^177^Lu decay model. This simulation provides organ-specific dose estimates based on an equivalent injected activity of the ^177^Lu-labeled conjugate. Individual organ cumulative and projected dose (Gy) is reported in table.

### No significant immunogenicity of vMET1-Fc in immunocompetent mice

To address the extent to which our non-mammalian antibodies may generate an immune response in mice, we performed a series of 7 weekly injections of vMET1-Fc, the commercial anti-MET antibody onartuzumab, or PBS, into C57BL/J6 mice at 1 mg/kg. To detect the anti-drug antibodies (ADAs), mouse plasma was collected and an ELISA was performed to check for mouse IgG binding to vMET1-Fc and onartuzumab. Our findings show that while plasma from mice injected with vMET1-Fc did give a higher ADA signal than mice injected with PBS (*P* = 0.01), the ELISA suggests that this is a comparable response to mice injected with onartuzumab (*P* = 0.54) (**Fig. 7A**). This suggests that a VNAR-Fc like vMET1-Fc will not generate an undue immune response, and, given the use of a human Fc, immunogenicity may be even be lower in a human system than in a murine system.

### Pharmacokinetics of vMET1-Fc in non-human primates

We next studied the pharmacokinetics of [^89^Zr]Zr-vMET1-Fc in two male rhesus macaques using PET/CT. Early scans at 1 h post-injection documented activity distribution in the circulatory system with no organ uptake and ROI analysis of several organs verified the activity uptake observed in PET/CT imaging (**Fig. 7B**, Supplementary Fig. S10). Clearance was predominantly observed through the kidneys, with an average SUV of 20.2 144 h post-injection (**Fig. 7C**). Remnants of first-pass clearance in the liver was also observed at this timepoint with an average standardized uptake value (SUV) of 6.3. Spleen, lung, and spinal cord ROIs observed minimal uptake of activity (< 2 SUV post injection 144 hours post injection) and no significant uptake of activity was observed in the brain or muscle of our primate models (< 1 SUV 144 hours post injection). By measuring the amount of decay-corrected activity in the blood at each time point, *in vivo* half-life was calculated to be 40.55 and 45.27 hours for each primate (**Fig. 7D**). Dosimetry estimates from injections of same initial activity of [^177^Lu]Lu-vMET1-Fc were performed, with the projected dose estimated to be 4 Gy to both liver and kidneys (**Fig. 7E**).

To evaluate the safety profile of vMET1-Fc, blood samples were drawn prior to each injection. Complete blood counts (CBC) and blood chemistry measurements were taken immediately. CBC and blood count analysis showed no concerning trends overall (**Supplementary Fig. S11 - S13**). We noted a transient increase in the liver enzyme aspartate aminotransferase 24 hours after our first injection, but these returned to baseline levels a day later.

## Discussion

Antibody-based therapies targeting MET-expressing NSCLC have demonstrated efficacy and are now used in the clinic. Telisotuzumab vedotin-tllv, a MET-targeting antibody–drug conjugate, is FDA-approved for previously treated NSCLC with high MET protein overexpression based on the LUMINOSITY trial, which reported a 35% objective response rate and a 7.2-month median duration of response (42). However, its utility is constrained by high MET express (≥ 50% of tumor cells with strong [3+] staining), modest durability, and treatment-related toxicities including peripheral neuropathy and edema. Similarly, the bispecific EGFR/MET antibody amivantamab has shown activity in METex14 disease via the CHRYSALIS study (43); yet, it remains unapproved for MET-expressing tumors, and its bispecific design may contribute to increased off-target effects like infusion-related reactions and dermatologic events (44). Earlier MET-targeting antibodies, including the monovalent onartuzumab and the bivalent emibetuzumab, failed to improve survival or demonstrate meaningful clinical benefit in late-phase trials (45–48). While these findings support the study of MET-directed antibodies as a viable therapeutic approach in NSCLC, the limitations of current agents underscore the need for next-generation antibody constructs with improved selectivity, durability, and broader therapeutic impact.

Towards this goal, we developed a shark-derived VNAR antibody that selectively targets the MET receptor as both an imaging and therapeutic agent. Our vMET1-Fc construct exhibited enhanced affinity compared to the monomer and demonstrated favorable *in vitro* properties, including selective, rapid, and sustained internalization into MET-expressing cells. *In vivo* studies revealed acceptable immunogenicity in murine models, in agreement with other findings of VNAR immunogenicity (49, 50). Importantly, Lu-177–labeled vMET1-Fc produced robust tumor growth inhibition in clinically relevant MET-amplified and METex14 NSCLC xenograft models, demonstrating its potential as a therapeutic agent. It is likely that the protruding CDR3 loop of vMET1 (Supplementary Fig. S1A) has allowed for a tighter engagement with the SEMA domain, as evidenced by the picomolar dissociation constant given by the bivalent Fc-based construct. The use of VNARs enhances developability; as a heavy-chain antibody, vMET1-Fc is simpler to produce and purify in large quantities, and humanization is now a well-established process, should it be required (51, 52).

While vMET1-Fc is an effective MET-binder, it exhibited no significant cytotoxicity and did not stimulate proliferation, which is a concern when targeting receptor tyrosine kinases (RTKs) (53, 54) that can necessitate approaches like monovalency (21). This should provide a lower selective pressure on the tumor environment and slow the development of resistance. The effective and selective internalization of vMET1-Fc into MET-expressing cells is of particular importance for the development of the construct into an effective PET imaging tracer or radiotherapeutic (55). Internalization allows the radionuclide a longer tenure at the target site, enabling higher intensity images over time, and yields more effective dose delivery from therapeutic radiometals. Our VNAR represents a promising candidate to address the lack of clinically approved PET imaging tracers and radiotherapeutics for MET-expressing lung cancer.

Antibody-based radionuclide therapy offers several distinct advantages over antibody–drug conjugates (ADCs) for targeting MET-expressing tumors. First, MET signaling directly regulates cellular responses to ionizing radiation, creating an intrinsic biological synergy that can be therapeutically exploited (56–61). Disruption of MET signaling in MET-overexpressing tumor cells impairs DNA damage repair following radiation-induced double-strand breaks, thereby enhancing cytotoxicity (62, 63). Consistent with this mechanism, inhibition of MET with clinically relevant small-molecule inhibitors such as savolitinib and capmatinib exhibits radiosensitizing activity in MET-amplified and MET exon 14–skipping NSCLC xenograft models (*39, 40*). Radiation-induced MET upregulation can establish a feed-forward mechanism that increases surface receptor density, improving antibody accessibility and promoting enhanced accumulation and retention of MET-targeted radioconjugates precisely when DNA damage response pathways are engaged. The resulting increase in intracellular radiation dose intensifies radiotoxic stress and counteracts MET-driven adaptive resistance. A precedent for this approach has been demonstrated in preclinical prostate cancer models, where targeted α-particle therapy induced DNA damage and antigen upregulation, thereby amplifying therapeutic efficacy (64).

A second advantage of radionuclide therapy is that the cytotoxic effects of radionuclide-emitted radiation are target-agnostic at the cellular level, enabling effective elimination of resistant tumor subclones and addressing intratumoral heterogeneity. Unlike ADCs, radionuclide therapies do not require receptor internalization, as emitted radiation exerts lethal effects through proximity and crossfire, allowing tumor cell killing even in the setting of heterogeneous or low MET expression.

Third, radionuclide therapy enables quantitative, noninvasive assessment of target engagement and treatment delivery—capabilities not available with ADCs. Many clinical trials stratify patients using immunohistochemistry assays that target epitopes distinct from those recognized by the therapeutic antibody, a practice that has been questioned and may contribute to reduced clinical response rates (*48*). In contrast, Zr-89–based PET imaging allows pre-therapeutic assessment of tumor uptake, enabling patient selection and personalization of administered activity through prescriptive dosimetry. Subsequent treatment with Lu-177 permits real-time monitoring of therapeutic delivery and biodistribution via SPECT, facilitating adaptive treatment strategies that are not feasible with non-radiolabeled ADCs.

Finally, antibody-based radionuclide therapy offers substantial versatility through the use of alternative radioisotopes. The successful conjugation of Lu-177 to vMET1-Fc supports the potential incorporation of other therapeutic radionuclides, including α-emitters such as Ac-223, enabling modulation of radiation quality and mechanism of action. This flexibility, combined with radiation-induced immune modulation and dose-dependent induction of immunogenic tumor cell death, positions MET-targeted radionuclide therapy as a highly adaptable and potentially superior therapeutic strategy compared with ADCs.

Demonstrating the favorable translational properties of our antibody, non-human primate studies showed no significant tissue uptake of [^89^Zr]Zr-vMET1-Fc outside of clearance organs. Uptake in clearance organs kidney and liver were consistent with uptake values previously reported from other antibodies (*65*). These findings likely reflect normal pharmacokinetic behavior and may also be influenced by the absence of tumor tissue, which would otherwise serve as a sink for radiotracer uptake. A limitation of this study is that we did not account for the potential presence of occult disease sites with high MET expression, which could alter systemic biodistribution. Nevertheless, estimated absorbed dose from our Lu-177 payload is well below the proposed renal dose constraint of 23 Gy for radionuclide therapy (*66*). Importantly, safety assessments in non-human primates and mice revealed no significant changes in hematologic parameters or serum chemistry following administration of either vMET1-Fc or [^89^Zr]Zr-vMET1-Fc, supporting the conclusion that cytotoxicity risk is primarily payload-dependent rather than antibody-driven. Accordingly, future structural optimization of vMET1-Fc should prioritize minimizing off-target uptake in normal human tissues while maintaining efficient systemic clearance.

In conclusion, this work constitutes the first reported example of a MET-targeting VNAR. We demonstrate that vMET1 is a highly selective VNAR for human MET, exhibiting negligible cytotoxicity, no effects on MET signaling, minimal immunogenicity, and high *in vivo* stability. Robust internalization of vMET1-Fc supports its potential for development as an antibody-drug conjugate or as an antibody-based radionuclide therapy. In MET-altered NSCLC xenograft models, vMET1-Fc showed high specificity for MET-expressing tumors and favorable pharmacokinetic properties, enabling effective imaging following radiolabeling with Zr-89 or Lu-177 and detection by PET/CT or SPECT, respectively. Moreover, [^177^Lu]Lu-vMET1-Fc delivered therapeutic relevant doses of radiation to MET-expressing tumors and effectively delayed tumor growth and prolonged overall survival. Collectively, these findings support the further development of this novel shark-derived VNAR as a theranostic agent for patients with MET-altered NSCLC.

## Acknowledgements

This project was supported by the Specialized Program of Research Excellence (SPORE) program, through the National Cancer Institute (NCI), grant P50CA278595, University of Wisconsin Carbone Cancer Center Support Grant (P30CA014520), donor funds from John Hallick and the MET Crusaders, NIH/NCI R01 CA237272 (A.M.L.), NIH/NCI R01 CA233562 (A.M.L.), a 2018 Prostate Cancer Foundation Challenge Award (A.M.L.), a Prostate Cancer Foundation Young Investigator Award (A.M.L.), and Andy North and Friends. The authors acknowledge the University of Wisconsin Small Animal Imaging & Radiotherapy Facility for supporting this work. The authors thank James P. Zacny for manuscript formatting and editing assistance.

