## Supplemental Information for "A MET-Targeted Variable New Antigen Receptor (VNAR) Theranostic for Non-Small Cell Lung Cancer"

[8]Department of Pathobiological Sciences. University of Wisconsin School of Veterinary Medicine, Madison, WI-53705.

[†]These authors contributed equally to this work.

**Supplementary Methods**

**Shark husbandry.** Juvenile male nurse sharks (*Ginglymostoma cirratum*) were housed in a round fiberglass tank with an operational volume of $18m^3$, operating as a recirculating aquaculture system with approximately two water turnovers per hour. Water salinity (32-35 ppt), temperature (27° C), ammonia content (<0.03 ppm), nitrogen dioxide (<0.1 ppm), nitrate (<0.1 ppm), pH (7.8-8.3), and dissolved oxygen (>5.0 mg/L) were monitored daily. Sharks were monitored on a weekly basis by veterinary staff for signs of illness or parasitism, and were fed at a rate of 5% body weight per week, split into three feedings. Diet consisted of octopus, Arctic char, red snapper, grouper, and shrimp.

**Immunization**. A shark weighing 3-4 kg was sedated via a 5-8-minute submersion in 0.01% tricaine mesylate (MS-222, Syndel, Ferndale, WA) solubilized in artificial seawater. Blood was drawn from the caudal vein using a 20G needle affixed to a syringe pre-filled with 1 mL of 1% EDTA. Blood (9 mL) was collected immediately prior to the first immunization injection, and subsequently every two weeks for 18 weeks. Initial immunization was performed with 200 ug of recombinant human MET (MET-H82E1, Acro Biosystems) emulsified 1:1 with Complete Freund's Adjuvant, which was then injected subcutaneously in the lateral fin. After one month, a booster injection, now with Incomplete Freund's Adjuvant, was similarly injected in the opposing lateral fin. Three further booster injections were administered at monthly intervals with declining amounts of MET protein (100, 100, 50 µg). Final three injections were given intravenously in the caudal vein, with antigen suspended in PBS supplemented with 350 mM urea and a final concentration of 500 mM NaCl. Plasma and buffy coat fractions were isolated from whole blood samples after centrifugal fractionation at 300 RCF for 5 min at 4° C with rotor breaks off. Immunization progress was monitored by screening plasma samples for the presence of MET-responsive IgNARs using biolayer interferometry.

**VNAR phage-display library construction.** Buffy coat samples giving a strong anti-MET BLI response were used for library assembly. RNA was isolated from samples using an RNeasy minikit (Qiagen, Hilden, Germany), and converted to cDNA via a High Capacity RNA-to-cDNA kit (Applied Biosystems, Thermo Fisher Scientific, Waltham, MA). Oligonucleotide primers adapted from Dooley, H. 2022(1), specific to the regions flanking frameworks 1 and 4 of IgNAR V regions, were used to specifically amplify VNAR-encoding cDNA using a polymerase chain reaction (PCR). PCR amplicons were separated by size using agarose gel electrophoresis, and nucleic acid bands of approximately 400 bp were excised, purified, restriction digested, and ligated into a pADL-22c phagemid vector (Antibody Design Labs, San Diego, CA). PCR and cDNA synthesis steps were each performed in 32 independent reactions, and then pooled after gel excision. Electrocompetent TG1 cells (Lucigen, Madison, WI) were electroporated (25 µF, 1500 V, 200 Ω) in six separate aliquots given 20 ng ligated DNA each. After 1-hour recovery in liquid recovery media, cells were plated on selective media dishes and incubated overnight at 30° C. Library titer was determined by serial dilution to be 1.9 x 108. After overnight growth, E. coli bacterial lawns were scraped and pooled, OD600 was measured. Pooled cells were and supplemented to 20% glycerol, aliquoted, and stored at -80° C. Phage were produced from E. coli stocks of the immunized anti-MET VNAR phagemid library using standard phage rescue methods and purified by precipitation with 4% (w/v) polyethylene glycol 8000 and 0.5 M NaCl. Titer of purified phage was then determined by serial dilution, infection of naïve TG1 cells, and plating on selective media plates.

**Phage-display biopanning.** Phage purified from the immunized anti-MET VNAR phage display library was used to screen for clones that bind to recombinant human MET. Recombinant MET was purchased biotinylated from the vendor. Biotinylated MET (10 ug) was exposed to Dynabeads M-270 Streptavidin (Invitrogen, Carlsbad, CA) in 2% (w/v) BSA in PBS. A volume of purified phage representing 1000-fold more particles than the size of the library was diluted in PBS to 1 mL. The protein-coated beads and phage were then combined in a microcentrifuge tube, and incubated 1 hour with end-over-end mixing. Beads were then washed 20 times with PBS and moved to a new tube to eliminate low-affinity binders. Phage bound to beads after washes was eluted with 1 mL of 100 mM triethylamine and neutralized with 500 µL of 1 M Tris-HCl, pH 7.5. TG1 cells were infected with neutralized phage and the process was repeated for a second round of biopanning. After the second round, SS320 E. coli cells were also infected for ELISA-based screening of VNARs.

**Production of VNAR monomer.** The DNA sequence encoding lead VNAR vMET1 was codon optimized for expression in E. coli systems and synthesized as a double stranded gBlock (Integrated DNA Technologies, Coralville, IA). This gBlock was cloned using restriction enzymes BamHI and HindIII (New England Biolabs (NEB), Ipswich, MA) into an in-house, pSUMO derived vector, featuring an 8x histidine and SUMO tag on the N-terminal end of the expressed region, separated from the VNAR by a TEV cleavage site. Cloned plasmid was then transformed into SHuffle T7 Competent E. coli K12 cells (NEB, Ipswich MA). Individual colonies were selected and amplified overnight at 37° C in 10 mL Terrific Broth with 50 µg/mL kanamycin and 0.04% glycerol. A sequence-confirmed colony was then expanded to 1 L and protein production was induced by adding 1 mL of 1 M IPTG, and incubating culture shaking overnight at 18° C. Cells were harvested by centrifugation at 5000 RCF for 15 minutes, and cells were lysed by sonication. Cell lysate was then spun for 30 minutes at 50,000 RCF, and the resulting supernatant sterile filtered through a 0.45 µm filter. VNARS were captured from this solution using a 5 mL HisTrap column (Cytiva, Marlborough, MA) at 5 mL/min flow rate. The resulting elution from this column was subjected to cleavage by TEV protease, from its 8x histidine and SUMO tag. Cleaved protein solution was again passed through a 5 mL HisTrap column, and the tagless flow-through was kept for further purification. This solution was passed through a HiLoad Superdex 75 size exclusion chromatography column (Cytiva, Marlborough, MA) using a mobile phase of PBS on an ÄKTApure fast protein liquid chromatography system (FPLC); chromatograms were obtained by monitoring UV absorbance at 280 nm. Fractions corresponding to a single peak of UV absorbance corresponding to the molecular weight of the vMET1 VNAR were collected and pooled. The resulting purified VNAR was then confirmed by size using agarose gel electrophoresis, and for binding to MET by BLI. Confirmed VNAR monomer was then diluted to 1 mg/mL in PBS, dispensed into 1 mL aliquots, flash frozen, and stored at -80° C.

**Production of VNAR-Fc.** The DNA sequence encoding lead VNAR vMET1 was codon optimized for expression in Chinese hamster (Cricetulus griseus) systems and synthesized as a double stranded gBlock (Integrated DNA Technologies, Coralville, IA). This gBlock was then integrated into a mammalian expression vector encoding the Fc domain of a human IgG1 (TGEX-SCblue, Antibody Design Labs, San Diego, CA), by homologous recombination, using an In-Fusion HD cloning kit (Takara Bio USA, San Jose, CA). Sequence confirmed plasmid encoding the VNAR-Fc was then transfected into ExpiCHO-S cells using an Expifectamine CHO Transfection Kit (Gibco, Thermo Fisher Scientific, Waltham, MA), per the manufacturer's recommendations. Cells were then cultured and maintained as recommended for 10-14 days, when cultures were harvested by centrifugation at 20,000 RCF for 30 minutes at 4° C. The resulting supernatant was supplemented with 300 mM NaCl and 20 mM Na2PO4, and passed through a 0.45 µm sterile filter. VNAR-Fcs were then captured by running the solution through a MabSelect PrismA column (Cytiva, Marlborough, MA) at 1 mL/min. Bound protein was then eluted using 100 mM citric acid, pH 3.0. Eluate was immediately neutralized with 2 M Tris-HCl, pH 9.0. The eluate was then purified further by size exclusion chromatography using a HiLoad Superdex 200 column on an ÄKTApure FPLC; chromatograms were obtained by monitoring UV absorbance at 280 nm. Fractions corresponding to a single peak of UV absorbance corresponding to the molecular weight of the vMET1 VNAR-Fc were collected and pooled. The resulting purified VNAR was then confirmed by size using agarose gel electrophoresis, and for binding to MET by BLI. Confirmed VNAR-Fc was then diluted to 1 mg/mL in PBS, dispensed into 1 mL aliquots, flash frozen, and stored at -80° C.

**Cell culture.** A panel of cell lines from non-small cell lung cancer (NSCLC; EBC-1, UW-Lung-21, A549, MGH 915-4, HCC827, NCI-H441, NCI-H1648, NCI-H1975, and NCI-H1993), gastric cancer (SNU-5), and head and neck squamous cell carcinoma (HNSCC; Detroit 562) was used to screen for MET expression. The breast cancer cell line T-47D, which has no MET expression, served as a negative control. EBC-1 (high MET amplification), UW-Lung-21 (MET exon 14 skipping mutation) and T-47D (no

MET) were selected for further in vitro and in vivo experiments. The source and culture conditions for all cell lines are detailed in Supplementary Table S1. All cells were maintained at 37 °C in a humidified atmosphere with 5% $CO_2$. Culture media were changed regularly, and cells were passaged at 70% confluence. All cell lines were authenticated by short tandem repeat profiling and routine mycoplasma testing was performed with MycoStrip (InvivoGen).

**Flow cytometry.** Single-cell suspension of cells was prepared by trypsinization. About 0.5 × $10^6$ cells were incubated with 100 pM, 1nM, or 100 nM vMET1-Fc for 30 min on ice. Then, the cells were washed and stained with 1.25 µg/mL PE-labeled goat anti-Human IgG Fc (eBioscience) for 30 min on ice in the dark. The cells were washed three times and resuspended in a flow cytometry staining buffer (2% FBS and 0.1% NaN3 in PBS). Data was collected with the Attune Flow Cytometer (Thermo Fisher), and at least 10,000 viable cells were gated and analyzed with FCS Express (Version 7.18).

**Western Blot analysis.** For the screening of MET expression by western blot, cells were plated in 10 cm dishes and harvested for the basal levels of MET, p-MET, and GAPDH. For the effect of MET signaling pathway analysis, cells were seeded in 10 cm dishes and grown to approximately 75% confluency before being treated with either a vehicle control (PBS), 1 nM, or 10 nM vMET1-Fc. Cells were harvested 4 or 24 h after treatment. Capmatinib (5 nM) was used as a positive control for the inhibition of MET signaling. For hepatocyte growth factor (HGF) stimulation experiment, A549 (low MET expression) was stimulated with 50 ng/ml HGF (R&D Systems, 294-HG-005/CF) for 30 min after overnight serum starvation. Ten nM vMET1-Fc was treated for 30 min before and after addition of HGF. Cell lysates were prepared as previously described (2,3). Equal amounts of protein were separated by sodium dodecyl sulfate polyacrylamide gel electrophoresis (SDS-PAGE), transferred to PVDF (polyvinylidene difluoride) membranes, and probed with specific primary antibodies. Target proteins were detected by chemiluminescence using horseradish peroxidase (HRP)-conjugated secondary antibodies (BioRad) and following incubation with ECL-HRP substrates (SuperSignal West Dura Extended Duration Substrates, Thermo Fisher). The specific antibodies and sources are listed in Supplementary Table S2. The blots were imaged on an Odyssey XF imaging system (LICORbio) and the quantification of the protein band signals was assessed by Image Studio (version 6.1, LICORbio).

**Bioconjugation and Radiolabeling of Zr-89.** vMET1-Fc was site-specifically labeled at the Fc-based glycosylation sites with deferoxamine (DFO) using a GlyClick DFO kit (Genovis), following the manufacturer's protocol, to give a consistent degree of labeling (DOL) of 2. Zirconium-89 was provided by the University of Wisconsin Medical Physics Department (Madison, Wi). [$^{89}$Zr]Zr-oxalate in 1.0 M oxalic acid was adjusted to pH 7.5 with 2.0 M HEPES. vMET1-Fc-DFO in PBS (pH 7.5) was added to [$^{89}$Zr]Zr-oxalate solution (80 µg/mCi) and incubated at 32° C, shaking at 250 rpm for 1 hour. The labeled product was purified using a size-exclusion PD-10 column preequilibrated with PBS buffer.

**Murine models.** All animal studies were conducted under a protocol approved by the University of Wisconsin Institutional Animal Care and Use Committee. Four-to-five-week-old female Hsd:athymic Nude-Foxn1nu mice (Envigo) were used for imaging and therapeutic experiments. Xenografts were established by injecting each cell line (1-2x $10^6$ cells/mL; 100 µL per mouse) in 1:1 mixture of Matrigel (Corning) subcutaneously above the right shoulder blade. Immunocompetent non-tumor bearing female ICR mice (Jackson Laboratory) were used for toxicity study of 300 µCi of [$^{177}$Lu]Lu-vMET1-Fc.

**Bioconjugation and radiolabeling of Lu-177.** vMET1-Fc was conjugated to the bifunctional chelator p-SCN-Bn-DOTA (Macrocyclics) to enable subsequent radiolabeling with Lu-177. Prior to conjugation, the vMET1-Fc solution (1-2 mg/mL in PBS) was adjusted to pH 8.4 using a freshly prepared 0.2 M Chelex-treated $Na_2CO_3$ buffer (pH 10.0) to maintain optimal conditions for isothiocyanate reactivity and minimize metal contamination. A 3-fold molar excess of p-SCN-Bn-DOTA (25 mg/mL in DMSO) was added slowly, 2 µL at a time, with continuous gentle agitation to prevent local concentration spikes and potential antibody precipitation. The reaction mixture was incubated at 37 °C for 90 minutes under light-protected conditions to facilitate the chelator's covalent attachment to primary amines on the antibody fragment via thiourea bond formation. Following conjugation, the mixture was subjected to ultrafiltration using Amicon Ultra-0.5 centrifugal filters (30 kDa MWCO, Merck Millipore) to remove excess unbound chelator. The conjugated antibody was washed repeatedly (≥3X) with Chelex-treated PBS (pH 7.2) to ensure complete buffer exchange and removal of unreacted reagents and metal ions. The final purified conjugate was collected, quantified, and stored at 4 °C for short-term use or at –20 °C for longer-term storage until radiolabeling. vMET1-Fc conjugated with p-SCN-Bn-DOTA were radiolabeled with Lu-177 by mixing the conjugated solution with Lu-177, previously buffered with sodium acetate buffer (NaOAc, 1 M, pH 5.5), at a target-specific activity of 100 µg/mCi. Reaction was carried out at 37 °C for one hour on a shaker at 350 – 500 RPM. Following incubation, radiolabeling efficiency and radiochemical purity were assessed using instant thin-layer chromatography (iTLC) on silica gel strips, developed with 50 mM EDTA (pH 5.5) as the mobile phase. Free Lu-177 migrates with the solvent front, while the radioligand remains at the origin. Radiochemical purity was quantified using a radio-TLC scanner. Next, the radioligand is purified on PD-10 desalting columns equilibrated and eluted with sterile phosphate-buffered saline (PBS, pH 7.4) to remove unbound Lu-177 and low-molecular-weight components. Radioligand is usually obtained with a radiolabeling yield ranging from 85–98%.

**Non-human primate husbandry.** Per Animal Welfare Act regulations, macaques living at the WNPRC are provided enclosures with at least 4.3, 6.0, or 8.0 sq. ft. of floor space depending on the weight of the animal, measure 30, 32, or 36 inches high, and contain a tubular PVC or stainless-steel perch to allow an animal to utilize the vertical dimensions of the enclosure. Each enclosure is also equipped with a horizontal or vertical sliding door, an automatic water lixit, and a stainless-steel feed hopper. Each single unit is also equipped with removable dividers so that multiple living configurations can be created out of a single row or bank of enclosures. These removable dividers allow animals to be paired or to be placed in even larger social groups to enhance psychological well-being and to inspire species-typical behavior (e.g., grooming, play, allo-parenting, etc.). Prior to each injection in this experiment, animals were anesthetized using up to 7 mg/kg ketamine and up to 0.03 mg/kg dexmedetomidine, both via intramuscular injection, which was reversed at the conclusion of each procedure by injecting up to 0.3 mg/kg atipamezole (IV or IM). A blood draw was performed after anesthesia, but before each antibody injection.

**AlphaFold modeling.** Predictive model of vMET1 VNAR was generated by inputting the VNAR's amino acid sequence into UCSF ChimeraX 1.9. The generated best model was displayed, colored by predicted local distance difference test (pLDDT). Display of side chains was turned on for residues within CDR1 and CDR3. A graph displaying predicted aligned error (PAE) was generated alongside the model.

**Supplemental Figures**

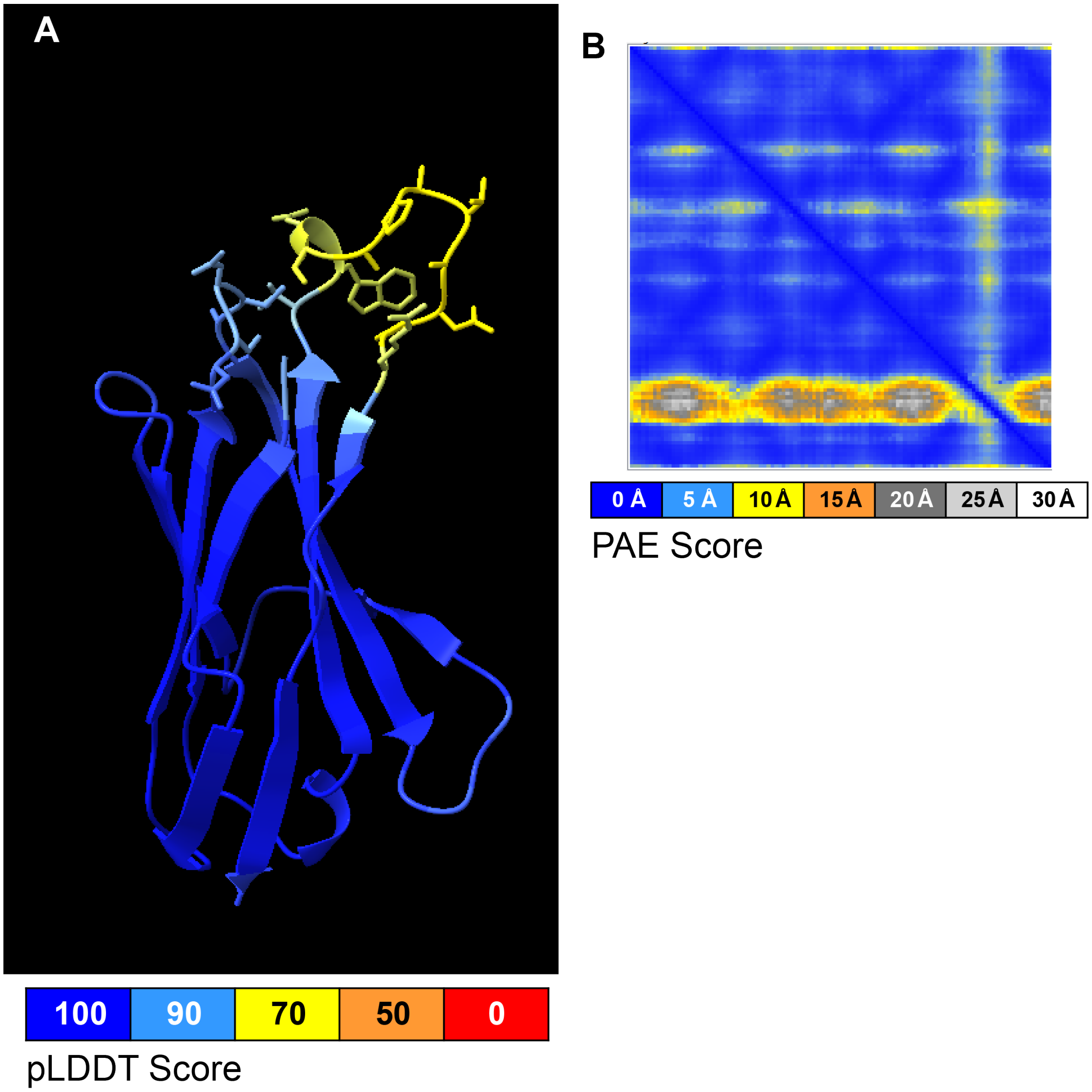


**C**

ARVDQTPQTITKETGESLTINCVLRDSNCALRSTYWYRKKSGSTREESICTGG
RYVETVNSGSKSFSLRINDLTVEDSGTYRCSAYETWDCAPATANPRAVYGRG
TAVTVN

**Supplementary Figure S1.** Alphafold predictive modeling of the vMET1 VNAR. **A,** The top predictive result of Alphafold modeling using ChimeraX software after inputting the sequence given in supplemental figure 1. Sequence is colored by the predicted local distance difference test (pLDDT), a per-residue measure of local confidence. Dark blue indicates a confidence near 100%, and a gradient follows to yellow, indicating a confidence near 70%. **B,** A predicted aligned area (PAE) graph of the structure given in **A**, which lists each residue along each axis, and shows a color at each (x, y) indicating expected position error at residue x if the predicted and true structures were aligned on residue y. **C,** The amino acid sequence of vMET1. CDRs are in red, and HV regions are in blue. Cysteine residues are highlighted; vMET1 is best described as a type II VNAR, although it has an extra cysteine in the HV2 region, resulting in five total cysteine residues.

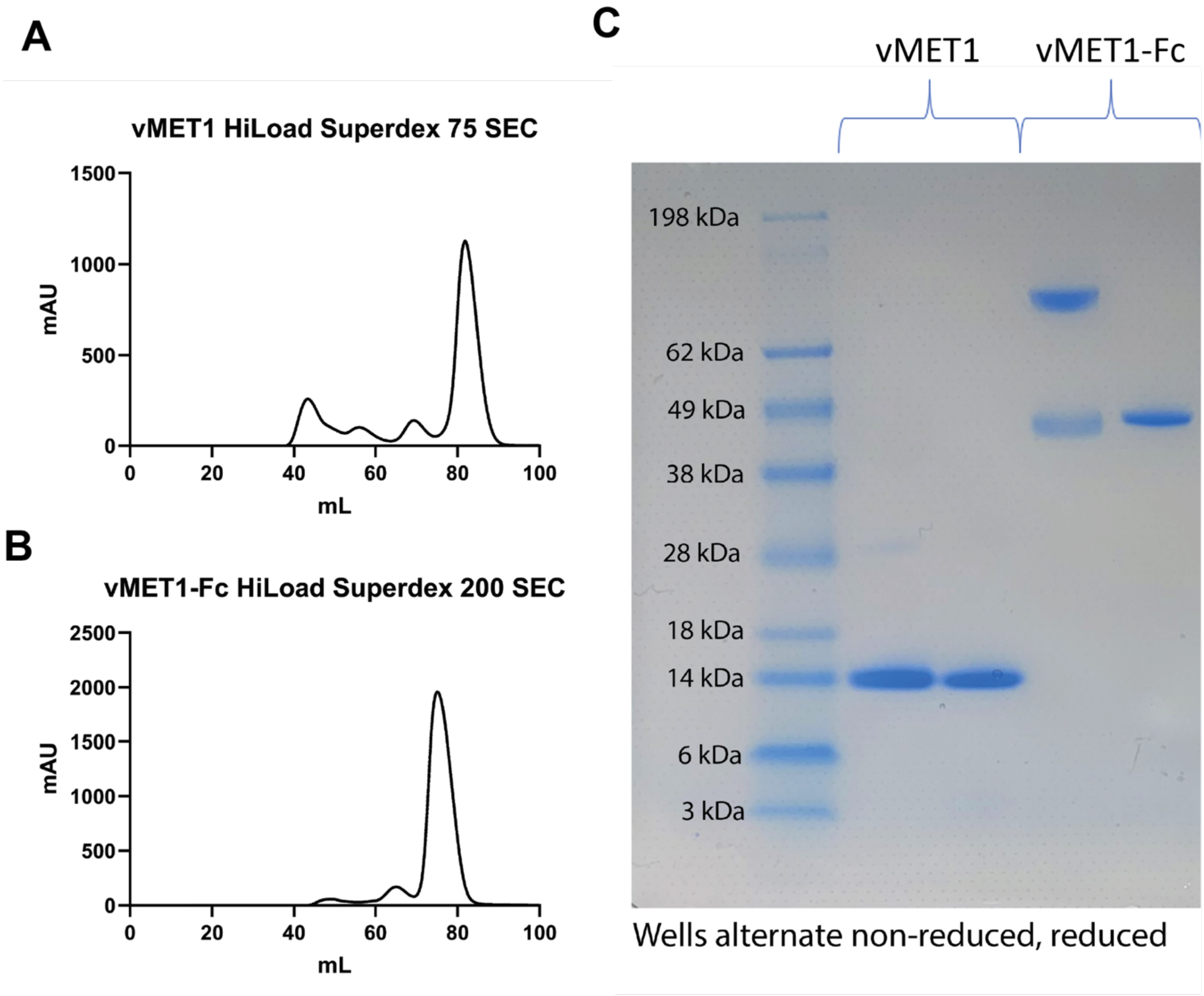


**Supplementary Figure S2.** vMET1 monomer and vMET1-Fc production and purification. **A,** A UV-measuring chromatogram at A280 showing the output of a HiLoad Superdex 75 size exclusion chromatography column after HisTrap purification of vMET1 monomer. The large peak on the right was taken for validation. **B,** A UV-measuring chromatogram at A280 showing the output of a HiLoad Superdex 200 size exclusion chromatography column after Protein A purification of vMET1-Fc. The large peak on the right was taken for validation. **C,** A Coomassie blue stained SDS-PAGE gel with a SeeBlue ladder, followed by the purified vMET1 monomer taken from the rightmost peak in (A), and then purified vMET1-Fc, taken from the rightmost peak in (**B**). Two lanes of each protein are run, with the right lane of each protein run under reducing conditions.

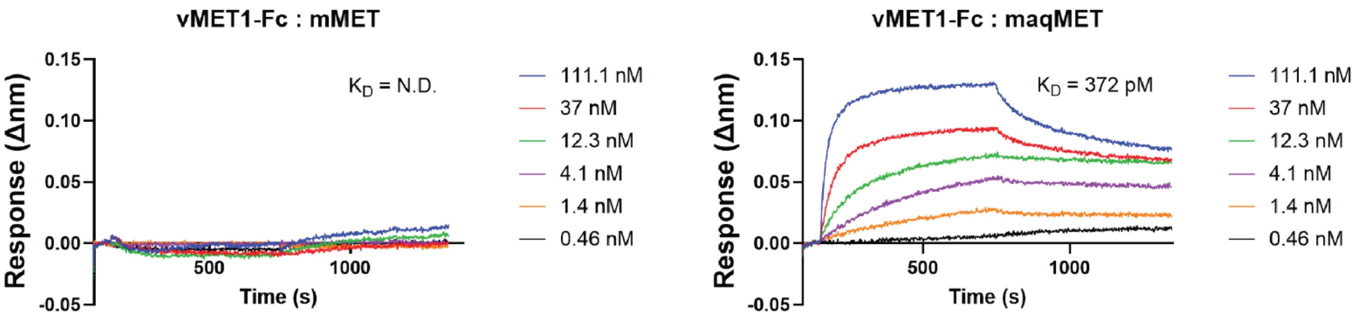


**Supplementary Figure S3.** vMET1-Fc species cross-reactivity assays. BLI sensorgrams of sensors loaded with **A,** murine MET or **B,** macaque MET exposed to serially diluted vMET1-Fc, followed by dissociation in assay buffer. Dissociation constants ($K_D$) are listed for each assay.

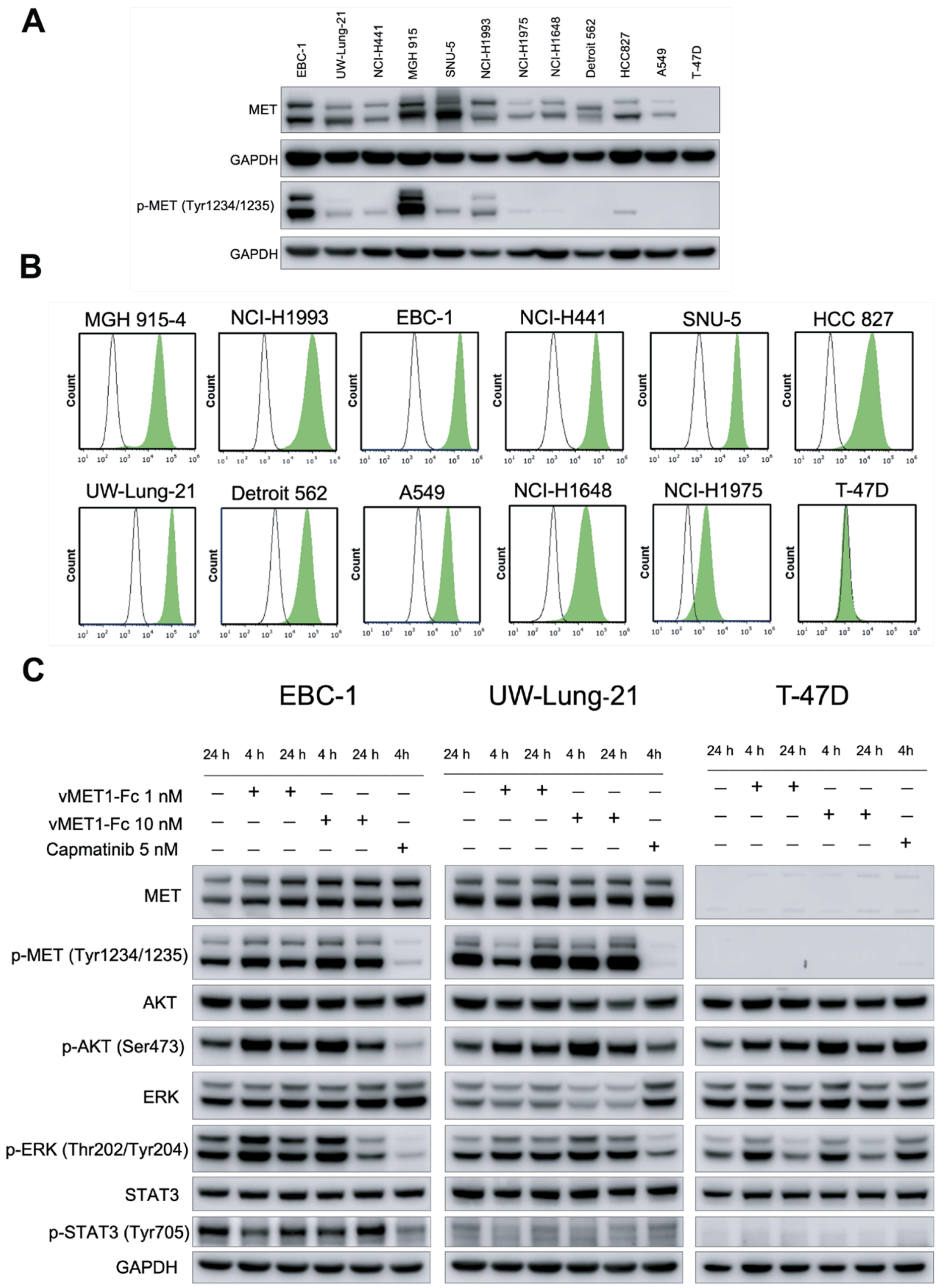


**Supplementary Figure S4.** vMET1-Fc does not inhibit downstream MET signaling in MET-altered Cell Lines. **A,** Western blot assaying levels of MET and p-MET in several cell lines. **B,** Flow cytometry results showing selective binding to MET cell lines that were exposed to 10 nM of vMET1-Fc , resulting in significant fluorescent signal (green) compared to their unexposed controls (grey). **C,** Western blots assessing MET, p-MET (Tyr1234/1235), AKT, p-AKT (Ser473), ERK, p-ERK (Thr202/Tyr204), and GAPDH in cells treated with vMET1-Fc demonstrated no effect on downstream signaling. Cells were treated with respective concentrations of vMET1-Fc for 4 h, and Capmatinib (5 nM) was used as a positive control.

A

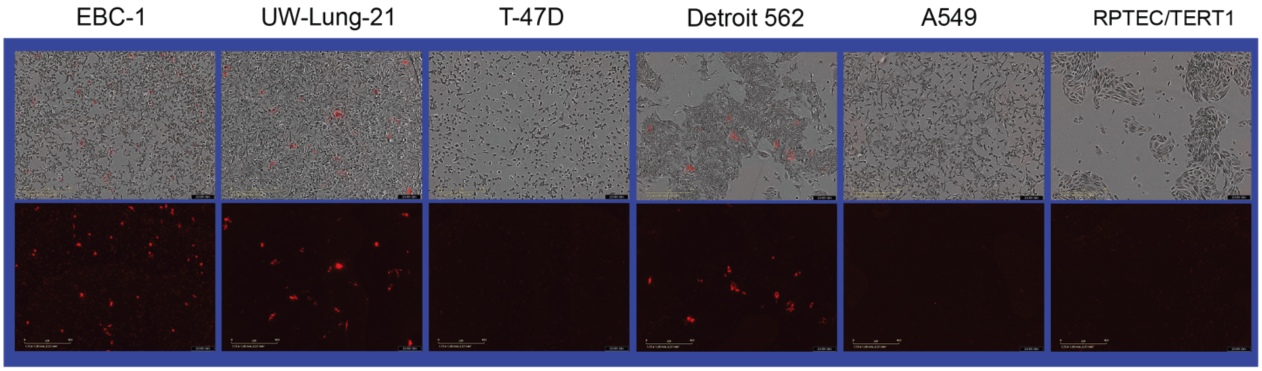


B

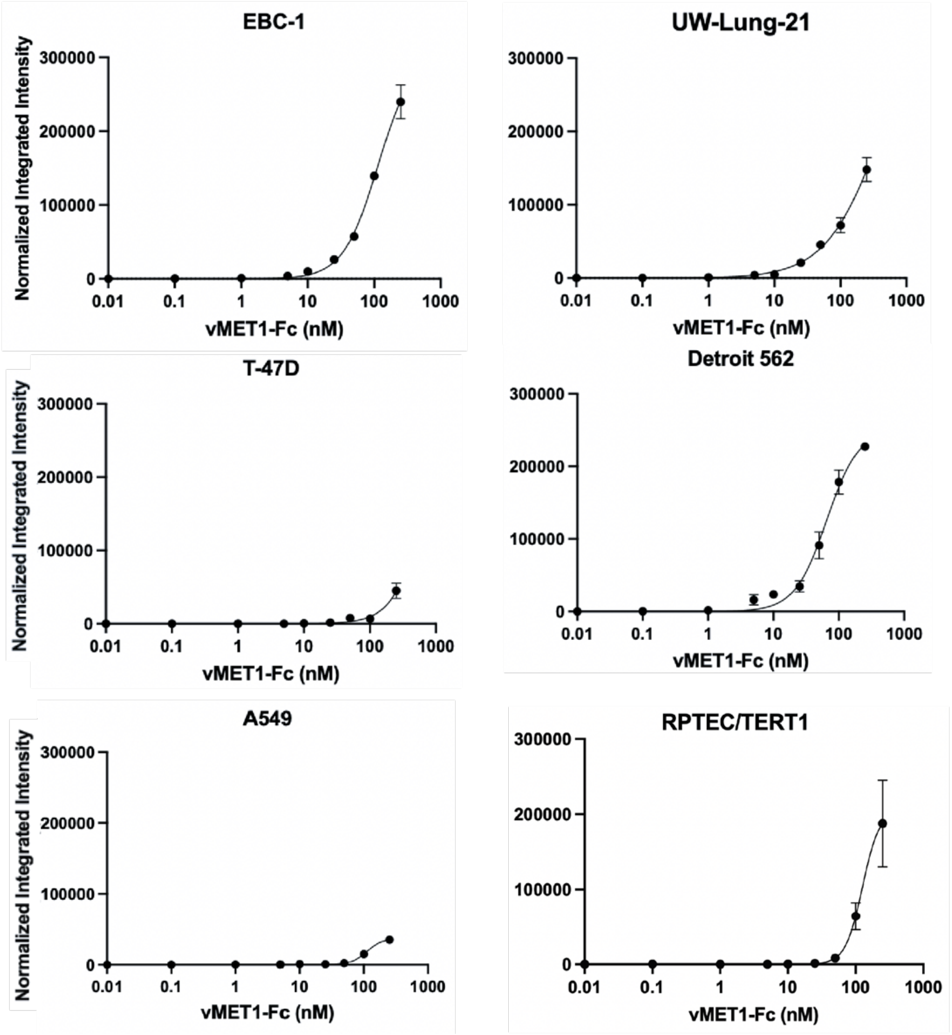


**Supplementary Figure S5.** vMET1-FC is primarily internalized by MET-expressing cells. **A,** Representative images from 10nM pHrodo Red internalization at 48 h post-treatment. Top row depicts an overlay of phase and orange imaging channels, while the bottom row shows orange imaging alone. **B,** Integrated fluorescent intensity calculated at 48 h after exposure to varying concentrations (0.1 – 250 nM) pHrodo Red labeled vMET1-Fc across cell lines.

**A**

EBC-1

| Mouse | Tumor Volume (mm³) | Tumor Diameter (mm) | Dose (Gy) | PVE Corrected Dose (Gy) |
|---|---|---|---|---|
| 1 | 591 | 10.41 | 17.65 | 18.54 |
| 2 | 502 | 9.86 | 15.65 | 16.65 |
| 3 | 683 | 10.93 | 11.66 | 12.10 |
| 4 | 495 | 9.82 | 24.09 | 25.64 |
| | | | **Mean** | 18.20 |
| | | | **SD** | 5.66 |

UW-Lung-21

| Mouse | Tumor Volume (mm³) | Tumor Diameter (mm) | Dose (Gy) | PVE Corrected Dose (Gy) |
|---|---|---|---|---|
| 1 | 344 | 8.69 | 16.43 | 18.09 |
| 3 | 44 | 9.47 | 19.54 | 20.98 |
| 4 | 158 | 6.70 | 24.53 | 29.42 |
| | | | **Mean** | 20.20 |
| | | | **SD** | 3.33 |

**B**

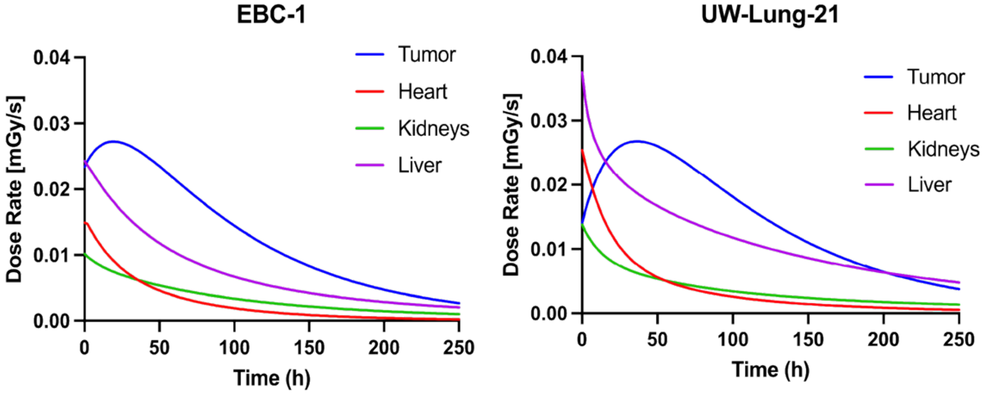


| Organ | Cumulative Dose (Gy) | Projected Dose (Gy) |
|---|---|---|
| Tumor | 9.25 | 12.51 |
| Heart | 1.99 | 2.17 |
| Kidneys | 2.08 | 4.00 |
| Liver | 4.56 | 11.5 |

| Organ | Cumulative Dose (Gy) | Projected Dose (Gy) |
|---|---|---|
| Tumor | 9.41 | 15.31 |
| Heart | 3.00 | 3.68 |
| Kidneys | 2.66 | 4.76 |
| Liver | 7.72 | 15.10 |

**Supplementary Figure S6.** Organ-based dosimetry from [$^{177}$Lu]Lu-vMET1-Fc shows effective tumor dose. **A,** Table of individual dose calculations per tumor Mean tumor volumes and absorbed dose estimates that received 300 µCi [$^{177}$Lu]Lu-vMET1-Fc calculated by RAPID Geant4 dosimetry platform, RAPID1. Partial volume effects (PVEs) and published specific absorbed fractions (SAFs) of 177Lu as a function of ROI volume, assuming a spherical tumor model, were also considered for the tumors. **B,** Dose rate profiles for tumor, kidneys, heart, and liver in EBC-1 and UW-Lung-21 mouse models estimated from IMALYTICS (bold lines are averages). Estimated dose is tabulated for each organ.

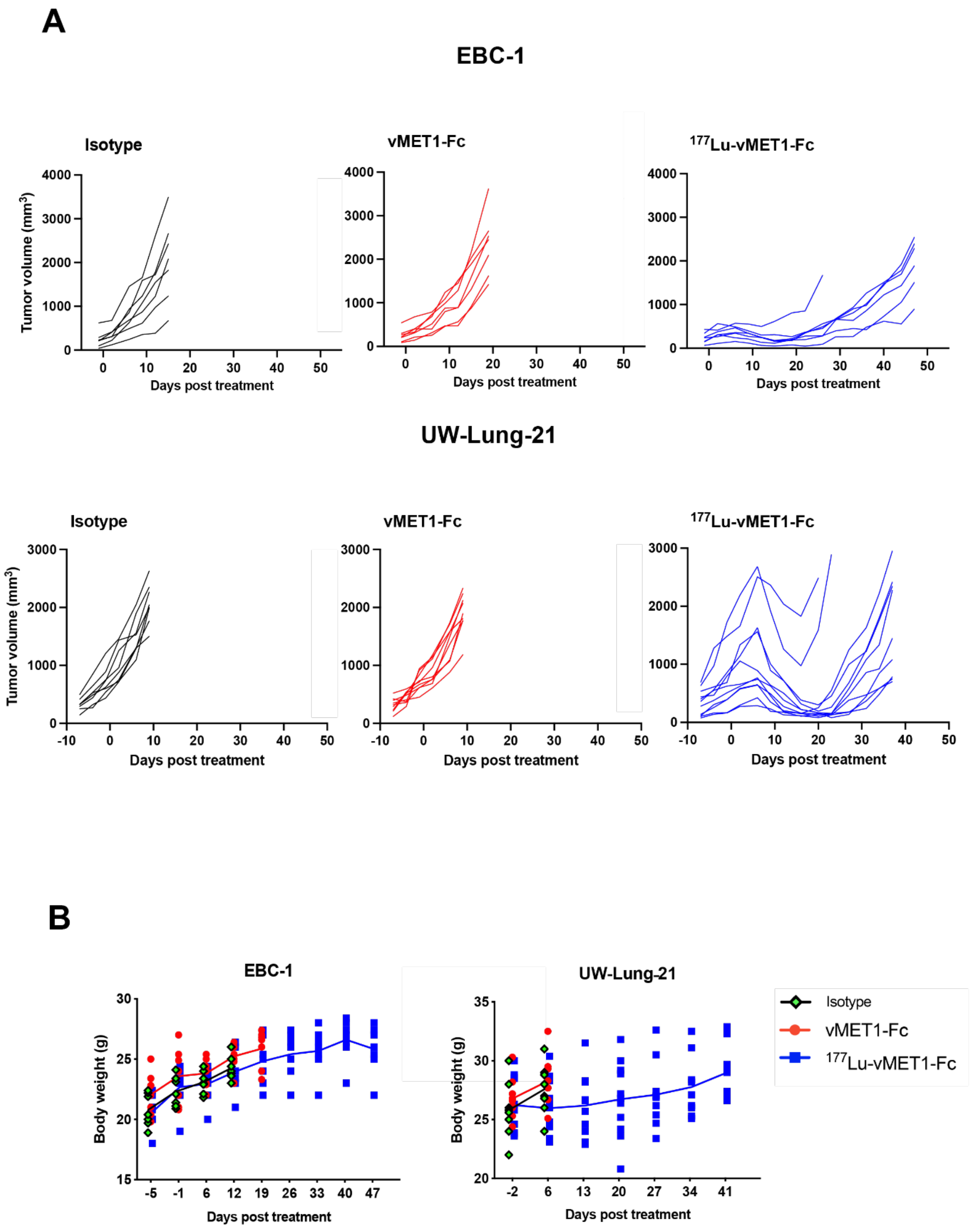


**Supplementary Figure S7.** [$^{177}$Lu]Lu-vMET1-Fc slowed individual tumor growth but did not strongly affect mouse body weight. **A,** Spaghetti plots of individual tumor volume measurements in mice that received isotype (1 mg/kg), vMET1-Fc (1 mg/kg), or [$^{177}$Lu]Lu-vMET1-Fc (300 μCi) in EBC-1 and UW-Lung-21. **B,** Serial measurements of total body weight revealed no significant differences in mass fluctuations between the various treatment cohorts in either EBC-1 or UW-Lung-21 tumor-bearing mice. Points are individual measurements; lines are trending averages per group.

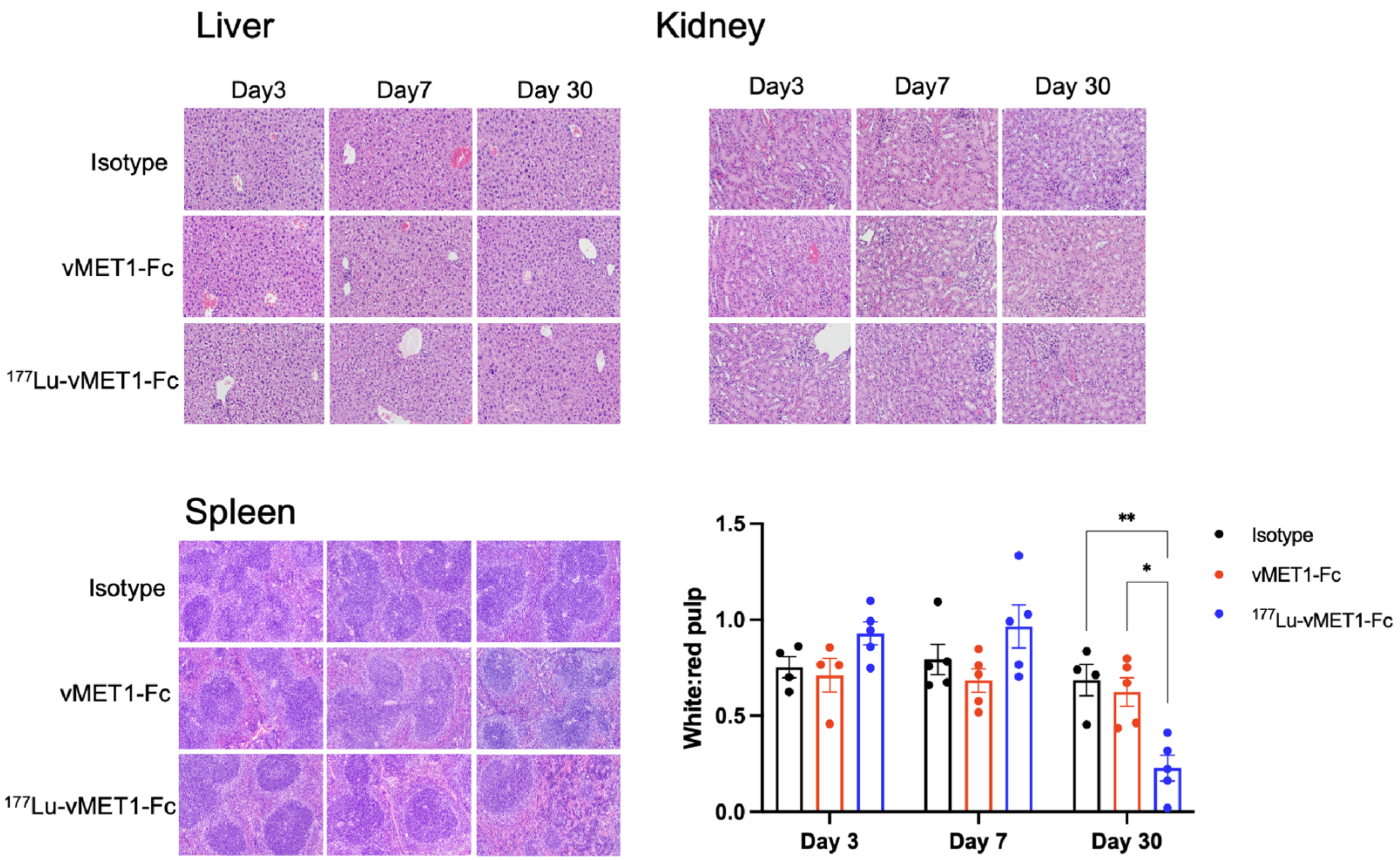


**Supplementary Figure S8.** Pathology screening for white and red pulp demonstrated expected reduction in white pulp content. H&E-stained sections of liver, kidney, and spleen collected at day 3, day 7, and day 30 are shown. Liver and kidney sections are included as representative histology. Spleen sections are shown together with quantification of white and red pulp using HALO Image Analysis software, presented as the white: red pulp ratio. Points are individual data points, *P < 0.05, ** P < 0.01

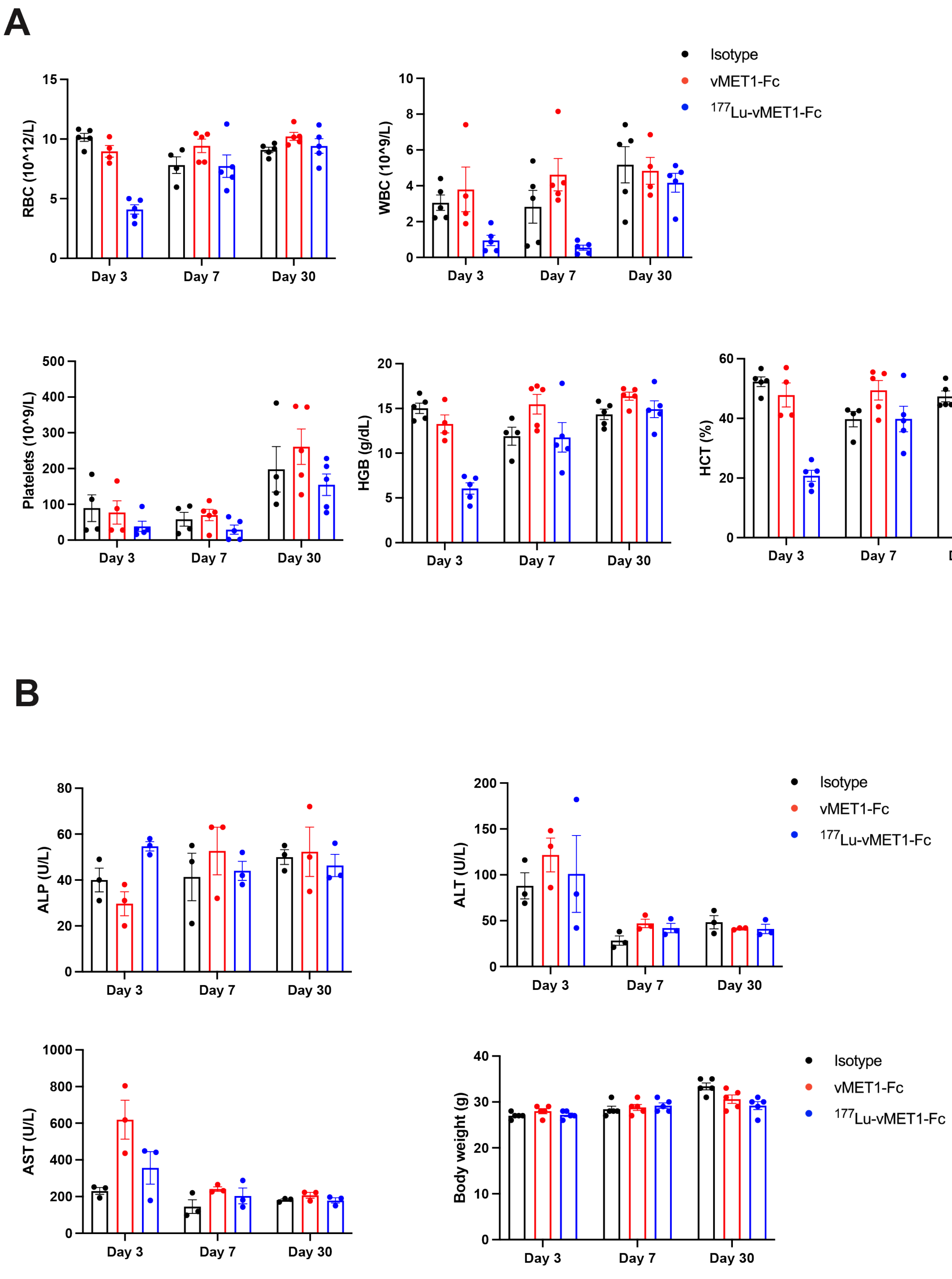


**Supplementary Figure S9.** [$^{177}$Lu]Lu-vMET1-Fc complete blood count (CBC) and blood chemistry analysis shows recoverable effect on blood cell population. **A,** Blood cell count and **B,** chemistry were performed on blood collected from mice on day 3 , day 7 and day 30 post-treatment (data points are individual values, bar graphs are means).

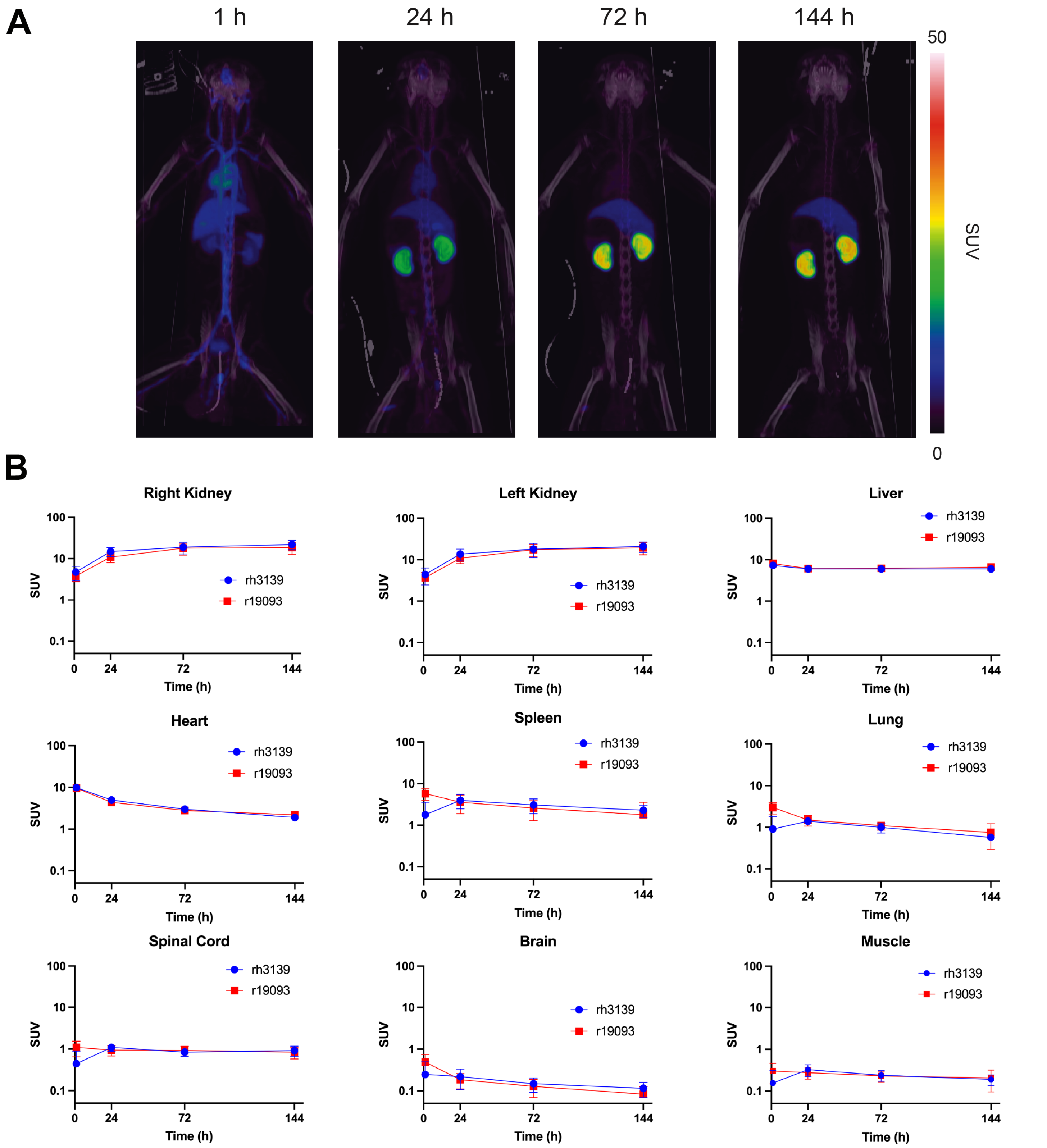


**Supplementary Figure S10.** Primate study shows expected clearance patterns. **A,** Representative PET/CT scan of second rhesus macaque not shown in Figure 7, at 1, 24, 72, and 144 h post-injection of 105.8 MBq (2.86 mCi) of [$^{89}$Zr]Zr-vMET1-Fc. **B,** Gamma counts of whole blood taken at each time point, with *in vivo* half-life calculated. **C,** ROI analysis of right and left kidney, liver, heart, spleen, lung, spinal cord, brain, and muscle for each primate quantifying uptake (points are average $SUV_{bw}$, bars are ROI SD). **D,** Estimated dose profile from [$^{177}$Lu]Lu-vMET1-Fc based on [$^{89}$Zr]Zr-vMET1-Fc uptake.

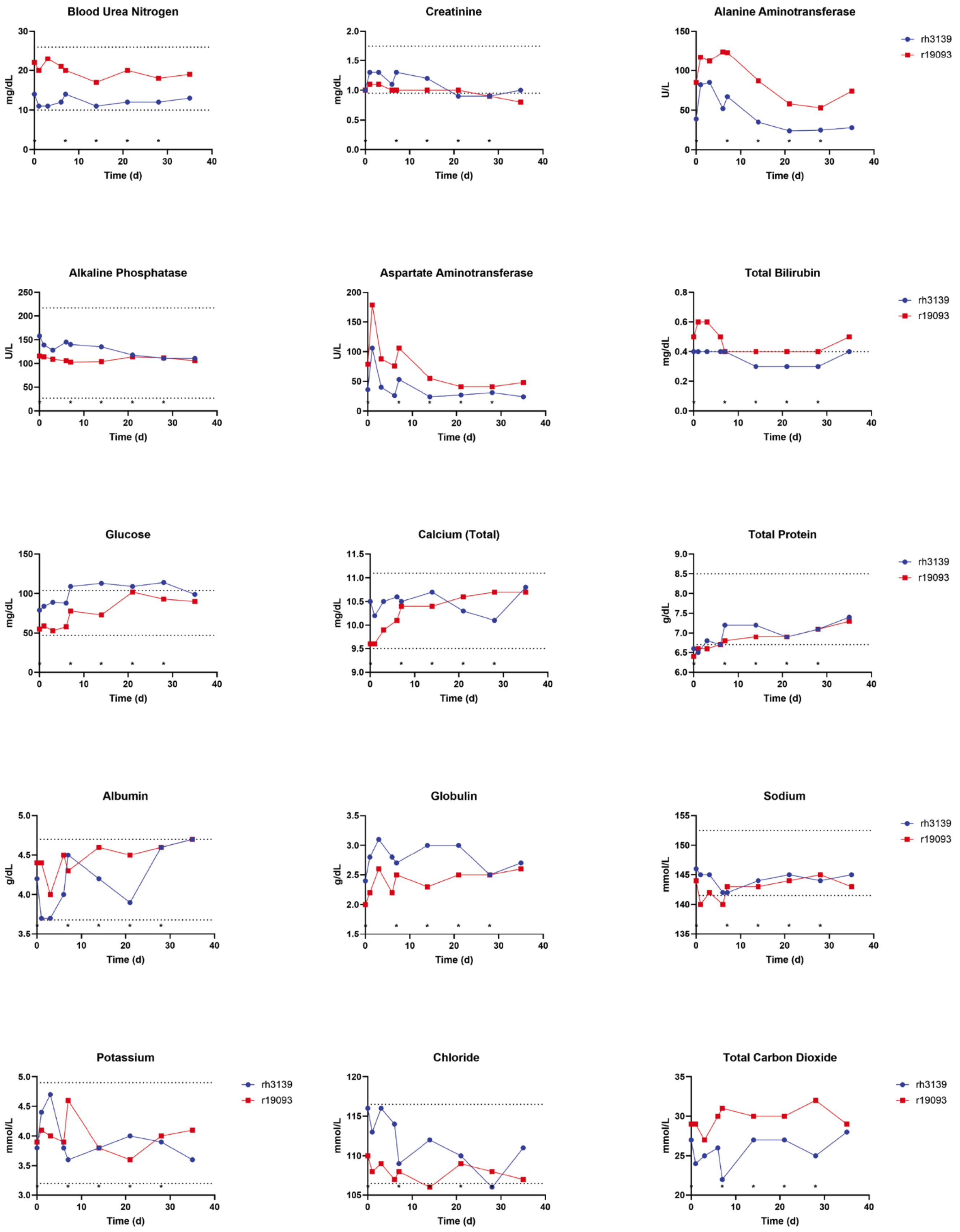


**Supplementary Figure S11.** Primate blood chemistry remains steady. Blood chemistry measurements of 15 metrics were taken for each primate at each time point and plotted against reference values where available.

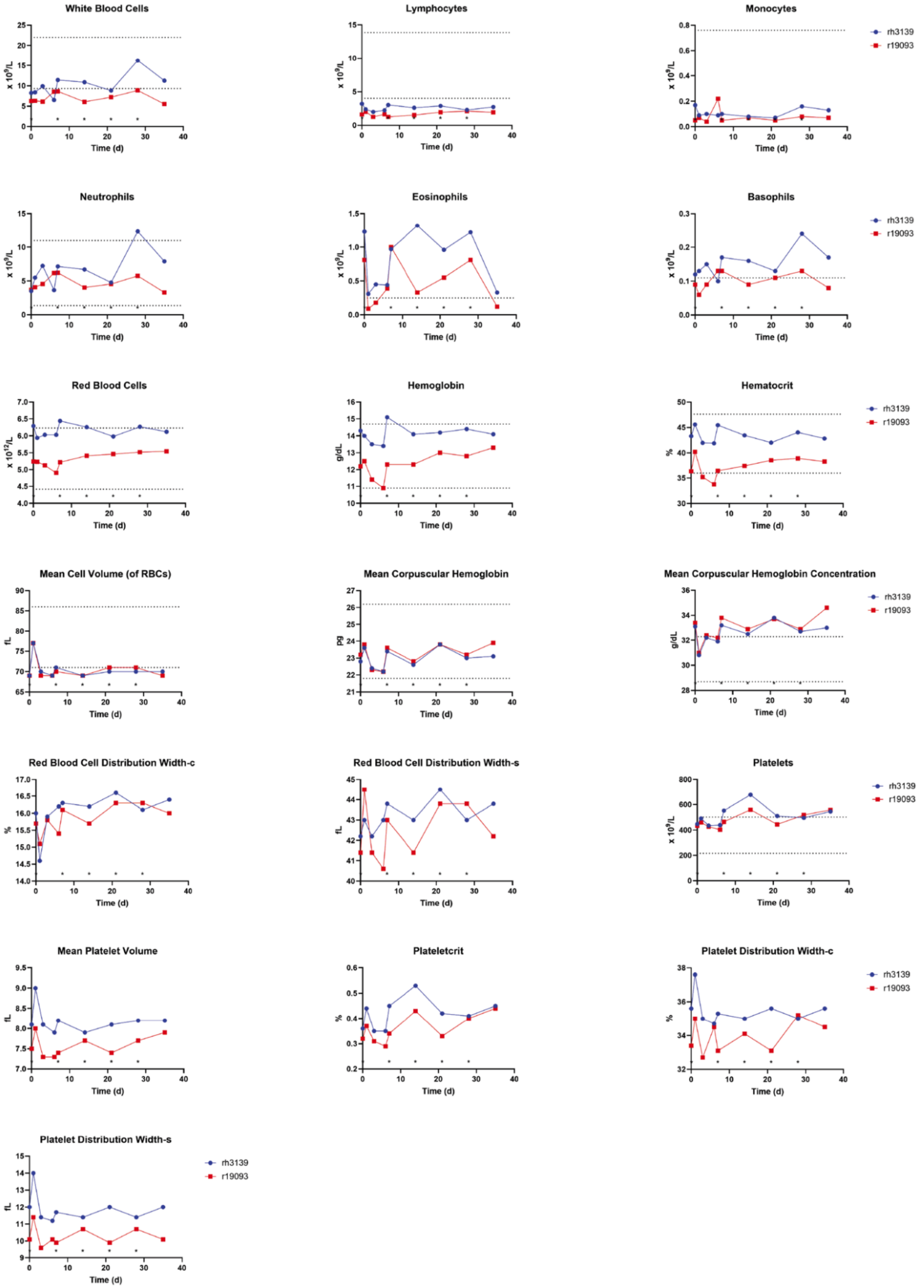


**Supplementary Figure S12.** Complete blood counts of primates show expected patterns. Blood cell counts of 19 metrics were taken from each primate at each time point and plotted against reference values where available.

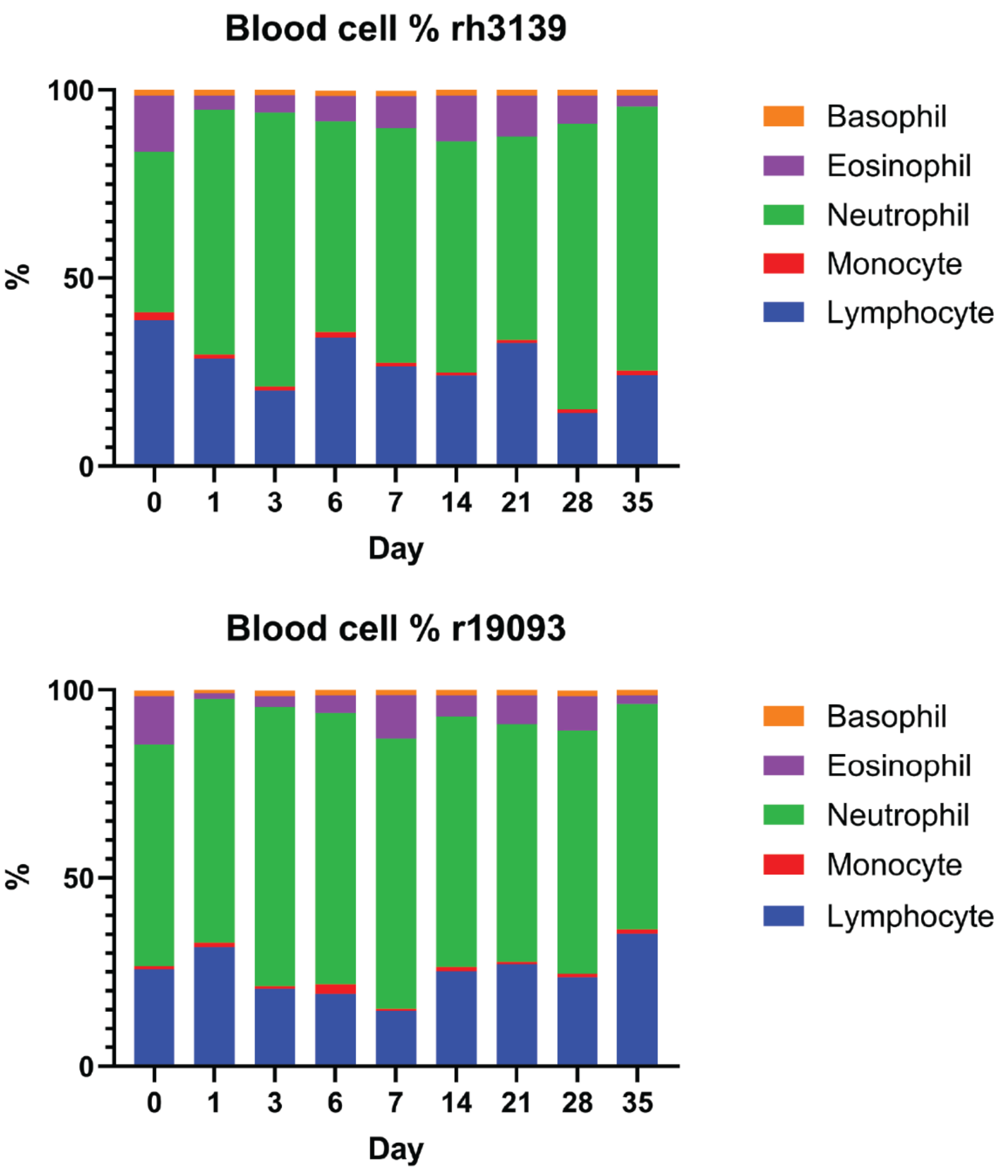


**Supplementary Figure S13.** Primate white blood cell ratios over time show no trending changes. Ratios were plotted for each primate after each blood draw throughout the 35 days of the study.

**Supplementary Table S1. Identified cell lines used in this paper with respective source and culture conditions.**

| Cell line | Source | Culture condition |
|---|---|---|
| EBC-1 | JCRB (Japanese Collection of Research Bioresources) Cell Bank Catalog #: JCRB0820 | EMEM, 200 mM L-glutamine, 10% FBS, penicillin (100 units/mL), streptomycin (100 mg/mL) |
| UW-Lung 21 | Established in Randall Kimple Lab (doi.org/10.1038/s41598-021-81832-1) | RPMI-1640, 10% FBS, penicillin (100 units/mL), streptomycin (100 mg/mL) |
| NCI-H441 | ATCC: HTB-174 | RPMI-1640, 10% FBS, penicillin (100 units/mL), streptomycin (100 mg/mL) |
| A549 | ATCC: CRM-CCL-185 | F-12K Medium, 10% FBS, penicillin (100 units/mL), streptomycin (100 mg/mL) |
| MGH 915-4 | Established in Aaron Hata lab (doi: 10.1158/1078-0432.CCR-19-3906) | RPMI-1640, 10% FBS, penicillin (100 units/mL), streptomycin (100 mg/mL) |
| NCI-H1648 | ATCC:CRL-5882 | RPMI-1640, 5% FBS, penicillin (100 units/mL), streptomycin (100 mg/mL) |
| NCI-H1975 | ATCC:CRL-5908 | RPMI-1640, 10% FBS, penicillin (100 units/mL), streptomycin (100 mg/mL) |
| NCI-H1993 | ATCC: CRL5909 | RPMI-1640, 10% FBS, penicillin (100 units/mL), streptomycin (100 mg/mL) |
| HCC827 | ATCC: CRL2868 | ATCC formulated RPMI-1640, 10% FBS, penicillin (100 units/mL), streptomycin (100 mg/mL) |
| SNU-5 | ATCC: CRL5973 | Iscove's Modified Dulbecco's Medium, 20% FBS, penicillin (100 units/mL), streptomycin (100 mg/mL) |
| T-47D | ATCC: HTB-133 | RPMI-1640, 10% FBS, 1 ml human recombinant insulin, penicillin (100 units/mL), streptomycin (100 mg/mL) |
| Detroit 562 | ATCC: CCL-138 | EMEM, 10% FBS, penicillin (100 units/mL), streptomycin (100 mg/mL) |
| RPTEC/TERT1 | ATCC: CRL-4031 | DMEM:F12 Medium, hTERT RPTEC Growth Kit, penicillin (100 units/mL), streptomycin (100 mg/mL) |

**Supplementary Table S2. Details regarding antibodies used in western blot analysis with their respective source, catalog number, supply company, and dilution ratio used in analysis.**

| Antibody | Source | Catalog # | Company | Dilution |
|---|---|---|---|---|
| MET | Rabbit | CST 8198 | Cell Signaling Technology | 1:1000 |
| phospho-Met (Tyr1234/1235) | Rabbit | CST 3077 | Cell Signaling Technology | 1:500 |
| AKT | Mouse | CST 2920 | Cell Signaling Technology | 1:1000 |
| phospho-AKT (Ser473) | Rabbit | CST 4060 | Cell Signaling Technology | 1:1000 |
| p44/42 MAPK (Erk1/2) | Mouse | CST 4696 | Cell Signaling Technology | 1:1000 |
| phospho-p44/42 MAPK (Erk1/2) (Thr202/Try204) | Rabbit | CST 4370 | Cell Signaling Technology | 1:500 |
| STAT3 | Mouse | SC-8019 | Santa Cruz Biotechnology | 1:1000 |
| phospho-STAT3 (Tyr705) | Rabbit | CST 9131 | Cell Signaling Technology | 1:500 |
| GAPDH | Rabbit | CST 5174 | Cell Signaling Technology | 1:1000 |
| HA-Peroxidase | Rat | 3F10 | Roche | 1:1000 |
| Mouse IgG-HRP | Goat | 31430 | Invitrogen | 1:10000 |
| Monkey IgG-HRP | Mouse | 4700-05 | Southern Biotech | 1:8000 |
| Onartuzumab | Human | A2468 | Selleck Chemicals | n/a |
